# Crossover interference and sex-specific genetic maps shape identical by descent sharing in close relatives

**DOI:** 10.1101/527655

**Authors:** Madison Caballero, Daniel N. Seidman, Jens Sannerud, Thomas D. Dyer, Donna M. Lehman, Joanne E. Curran, Ravindranath Duggirala, John Blangero, Shai Carmi, Amy L. Williams

**Affiliations:** Department of Molecular Biology and Genetics, Cornell University, Ithaca, NY 14853, USA; Department of Computational Biology, Cornell University, Ithaca, NY 14853, USA; South Texas Diabetes and Obesity Institute and Department of Human Genetics, University of Texas Rio Grande Valley School of Medicine, Brownsville, TX 78520, USA; Department of Medicine, University of Texas Health Science Center at San Antonio, San Antonio, Texas 78229, USA; Braun School of Public Health and Community Medicine, The Hebrew University of Jerusalem, Jerusalem, Israel

## Abstract

Simulations of close relatives and identical by descent (IBD) segments are common in genetic studies, yet most past efforts have utilized sex averaged genetic maps and ignored crossover interference, thus omitting features known to affect the breakpoints of IBD segments. We developed Ped-sim, a method for simulating relatives that can utilize either sex-specific or sex averaged genetic maps and also either a model of crossover interference or the traditional Poisson model for inter-crossover distances. To characterize the impact of previously ignored mechanisms, we simulated data for all four combinations of these factors. We found that modeling crossover interference decreases the standard deviation of the IBD proportion by 10.4% on average in full siblings through second cousins. By contrast, sex-specific maps increase this standard deviation by 4.2% on average, and also impact the number of segments relatives share. Most notably, using sex-specific maps, the number of segments half-siblings share is bimodal; and when combined with interference modeling, the probability that sixth cousins have non-zero IBD ranges from 9.0 to 13.1%, depending on the sexes of the individuals through which they are related. We present new analytical results for the distributions of IBD segments under these models and show they match results from simulations. Finally, we compared IBD sharing rates between simulated and real relatives and find that the combination of sex-specific maps and interference modeling most accurately captures IBD rates in real data. Ped-sim is open source and available from https://github.com/williamslab/ped-sim.

**Author summary:** Simulations are ubiquitous throughout statistical genetics in order to generate data with known properties, enabling tests of inference methods and analyses of real world processes in settings where experimental data are challenging to collect. Simulating genetic data for relatives in a pedigree requires the synthesis of chromosomes parents transmit to their children. These chromosomes form as a mosaic of a given parent’s two chromosomes, with the location of switches between the two parental chromosomes known as crossovers. Detailed information about crossover generation based on real data from humans now exists, including the fact that men and women have overall different rates (women produce ~1.6 times more crossovers) and that real crossovers are subject to *interference*—whereby crossovers are further apart from one another than expected under a model that selects their locations randomly. Our new method, Ped-sim, can simulate pedigree data using these less commonly modeled crossover features, and we used it to evaluate the importance of sex-specific rates and interference in real data. These comparisons show that both factors shape the amount of DNA two relatives share identically, and that their inclusion in models of crossover better fit data from real relatives.

## Introduction

Inferring identical by descent (IBD) segments and estimating relatedness are classical problems in human genetics [1], with recent work motivated by the abundance of close relatives in large samples [2–6]. In order to study individuals with a known relationship, many investigators have performed simulations, both to evaluate novel methods [4–8], and to characterize the properties of IBD sharing rates among relatives [9, 10]. Additionally, direct-to-consumer genetic testing companies—now with data from several million individuals— rely on simulated data to infer relationships by matching relatedness statistics from their customers to those from simulations [11, 12].

In parallel with the above, efforts to characterize crossovers, including the dynamics of crossover interference [13–15] and differences in male and female genetic maps [15–17] have yielded precise resources for realistically simulating this form of recombination. Despite this, most prior simulations and canonical models of IBD sharing between relatives [18] make use of sex averaged genetic maps and have ignored crossover interference.

Differences between male and female crossover rates were first identified decades ago [19], and modern data from families enable detection of separate male and female crossovers and therefore the inference of distinct maps [15–17]. Other commonly used but inherently sex averaged genetic maps [20] are based on population linkage disequilibrium (LD) patterns—the signal left by thousands of male and female meioses. Another source of crossover events is switches between ancestral populations within an admixed individual’s genome, but lacking information about which ancestor produced each crossover, the resulting admixture-based maps are also sex averaged [21]. Lastly, recent work on sequencing and resolving crossovers in single oocytes [22, 23] and sperm cells [24–27] provide information on sex-specific crossover properties and can be used to construct sex-specific maps.

In turn, characterization of crossover interference, initially observed as unexpectedly low rates of double crossovers in early *Drosophila* linkage analyses [28], now includes sex-specific parameter estimates from over 18,0 human meioses [15]. During meiosis, crossovers appear as chiasmata that physically link homologous chromosomes within tetrads—four chromosome bundles that consist of two copies of each homologous chromosome. Since chiasmata involve two non-sister chromatids (i.e., chromosome copies of distinct parental origin) the resulting crossover affects just two of the four gametes. To account for crossover interference, early models assumed that crossover intermediates are placed uniformly at random, but that only one of m intermediates resolves as a chiasma [29, 30]. The inter-chiasmata distance under this model is gamma distributed with an integer shape parameter *m* (the *χ*^2^ model). This and current models assume a uniformly random placement of chiasmata among the chromatids in a tetrad (i.e., no chromatid interference [22, 23]), corresponding to an independent probability of 1/2 for a gamete to contain any crossover. Later work found that inter-chiasmata distances are better fit by a gamma distribution with a fractional shape parameter [13] (the gamma model). Building on this, Housworth and Stahl found better fits to human inter-crossover distances using a mixture (two-pathway) model that, in addition to the gamma model, also includes some fraction of events that escape interference [14].

Here, we employ empirical human genetic maps and interference estimates to analyze the effects of crossover modeling on IBD distributions between close relatives. Specifically, we simulated several types of close relatives using either sex-specific or sex averaged crossover genetic maps [16], and either incorporating crossover interference [14, 15] or using a non-interference (i.e., Poisson) model. While mean IBD sharing rates are unaffected by these factors, the variance in IBD sharing proportion differs substantially between them, to a degree that impacts relationship classification metrics and estimates of the time since admixture in simulated admixed individuals. Furthermore, by analytically solving a theoretical renewal process model, we show that crossover interference also impacts the distribution of IBD segment lengths, and we confirm these results using simulations.

To determine which crossover model best fits data from real relatives, we leveraged genotypes from the San Antonio Mexican American Family Studies (SAMAFS) [31–33], a dataset comprising roughly 2,500 samples in dozens of pedigrees. With thousands of close relative pairs, these data enable precise estimates of IBD summary statistics. We also leveraged IBD sharing rates from 20,240 full sibling pairs analyzed by Hemani *et al*. [34]. We found that use of sex-specific genetic maps and interference modeling provide overall better fits to IBD sharing summary statistics in these real relatives than do other crossover models.

We conducted all simulations for this study using Ped-sim, an open source method we developed that performs simulations of relatives using either sex-specific or sex averaged genetic maps and either a model of crossover interference (the Housworth-Stahl model) or the traditional Poisson model (Methods).

## Results

We used Ped-sim to simulate 10,000 pairs of relatives for several relationship types and each of four crossover models. Ped-sim can produce genetic data for relatives given input haplotypes, but the analyses we present leverage exact IBD segments as detected through internally tracked haplotype segments (Methods). These segments arise by (simulated) descent from the chromosomes of founders—i.e., pedigree members whose ancestors Ped-sim does not model.

Comparing IBD sharing between simulated and real relatives is complicated by the fact that deeper, “background” relatedness from cryptic common ancestors can exist between real samples. This may inflate the relatedness between real samples above that implied by the more recent common ancestors that we focus on. Moreover, population-based IBD inference procedures are subject to both false positive and false negative signals, whereas simulated data perfectly capture the IBD generated under a given model.

We detected IBD in the real SAMAFS data using a family-based method [35] for the full siblings and a population-based approach [36] for other relatives. Similarly, the 20,240 Hemani *et al*. full sibling IBD estimates (Hemani20k) are from a family-based algorithm [34]. Family-based IBD detection largely obviates issues of background relatedness and false signals because it models haplotype transmissions from parents to children (in the case of the SAMAFS full siblings, also using data from the parents; Methods). We therefore consider the analyses of IBD sharing in full siblings as directly parallel to the simulated data.

The analyses of non-sibling relative types require adjustment, and we mean-shift the shared IBD proportion of each real data relationship class to match theoretical expectations. This does not address effects of imperfect IBD inference on the standard deviation of IBD sharing, which derives from squared deviations from the mean, and we consider the standard deviation quantities for real relatives to be biased except in the case of full siblings. For other relationships, we instead focus on quartiles of the IBD sharing fraction.

The IBD proportions quoted hereafter are fractions of the diploid genome two samples share, and we calculated them as half the fraction of the genome shared IBD1 plus the IBD2 fraction (Methods). The numerical IBD1/IBD2 types indicate how many haplotypes the pair shares IBD at a given location, one or two, respectively. Below, we abbreviate sex-specific and sex averaged as SS and SA, respectively, and refer to the four crossover models we used with Ped-sim as: SS+intf for sex-specific genetic map with interference; SS+Poiss for sex-specific map, Poisson event distribution (i.e., no interference); SA+intf for sex averaged genetic map with interference; and SA+Poiss for sex averaged map, Poisson event distribution.

### Sex-specific maps and interference oppositely affect variance in IBD sharing proportion

We simulated full siblings, first cousins, first cousins once removed, and second cousins under all four crossover models. For all relative types, use of SS genetic maps increases the variance in IBD proportion compared to the SA map, though the effect is somewhat limited. In particular, averaged among these relationships, the standard deviation increases by 3.6% under the Poisson crossover localization model and 4.7% under the interference model (Fig 1, S1). SS maps have similar effects on the size of the interquartile range, increasing this span by an average of 3.9% under the Poisson model and 5.3% in the presence of crossover interference. These small differences in IBD sharing statistics correspond to distributions of IBD rates for SS and SA maps that are difficult to distinguish visually (S2 Fig).

**Figure 1:**
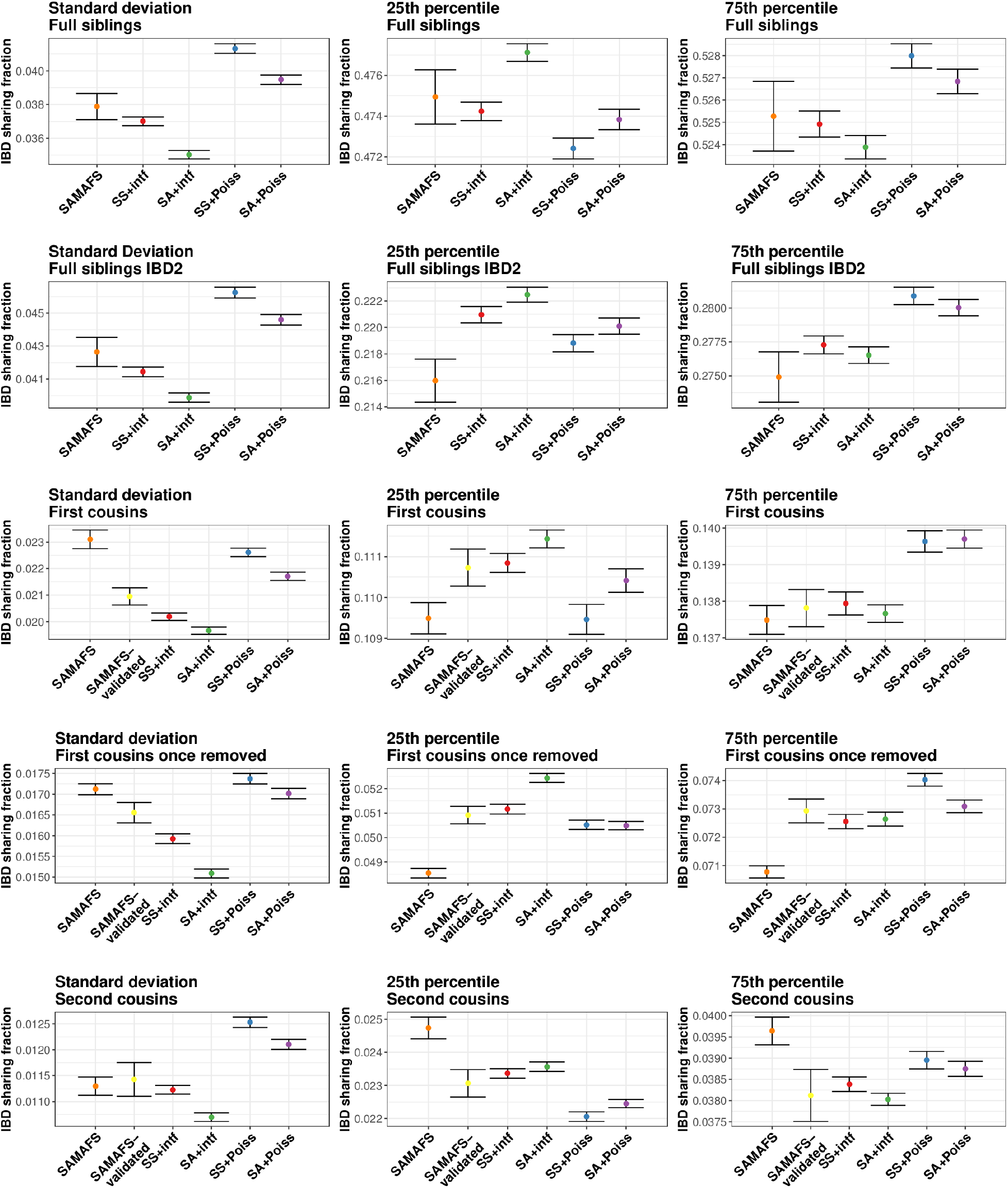
Standard deviations, 25th, and 75th percentiles of IBD sharing fraction in real and simulated data for full siblings through second cousins. Points are from the SAMAFS, SAMAFS-validated subset (except full siblings), Hemani20k set (only full siblings), and the simulation models. The latter are labeled using abbreviations given in the main text. The SAMAFS and SAMAFS-validated 25th and 75th percentiles are mean-shifted for the first cousins, first cousins once removed, and second cousins, but are unaltered for the full sibling and the full sibling IBD2 quantities. Bars indicate one standard error as calculated from 1,000 bootstrap samples.

By contrast, crossover interference has a strong effect on the variance in IBD sharing fraction, decreasing the standard deviation compared to the Poisson model by 10.0% when simulating with SS maps and 10.9% using the SA map (averaged over all relationships we considered; Fig 1). Furthermore, interference tightens the range between the 25th and 75th percentiles by 9.8% when using SS maps and 11.0% using the SA map. With decreased variances of these magnitudes, the distributions of IBD proportions for relatives simulated under interference are noticeably more peaked near the mean, with smaller tails (Fig 2). These results highlight the importance of including interference when simulating relatives, and hint that distantly related samples may have non-zero IBD sharing more frequently when simulated under interference—a feature we analyze below (see “Rates of sharing at least one IBD segment among distant relatives”).

**Figure 2:**
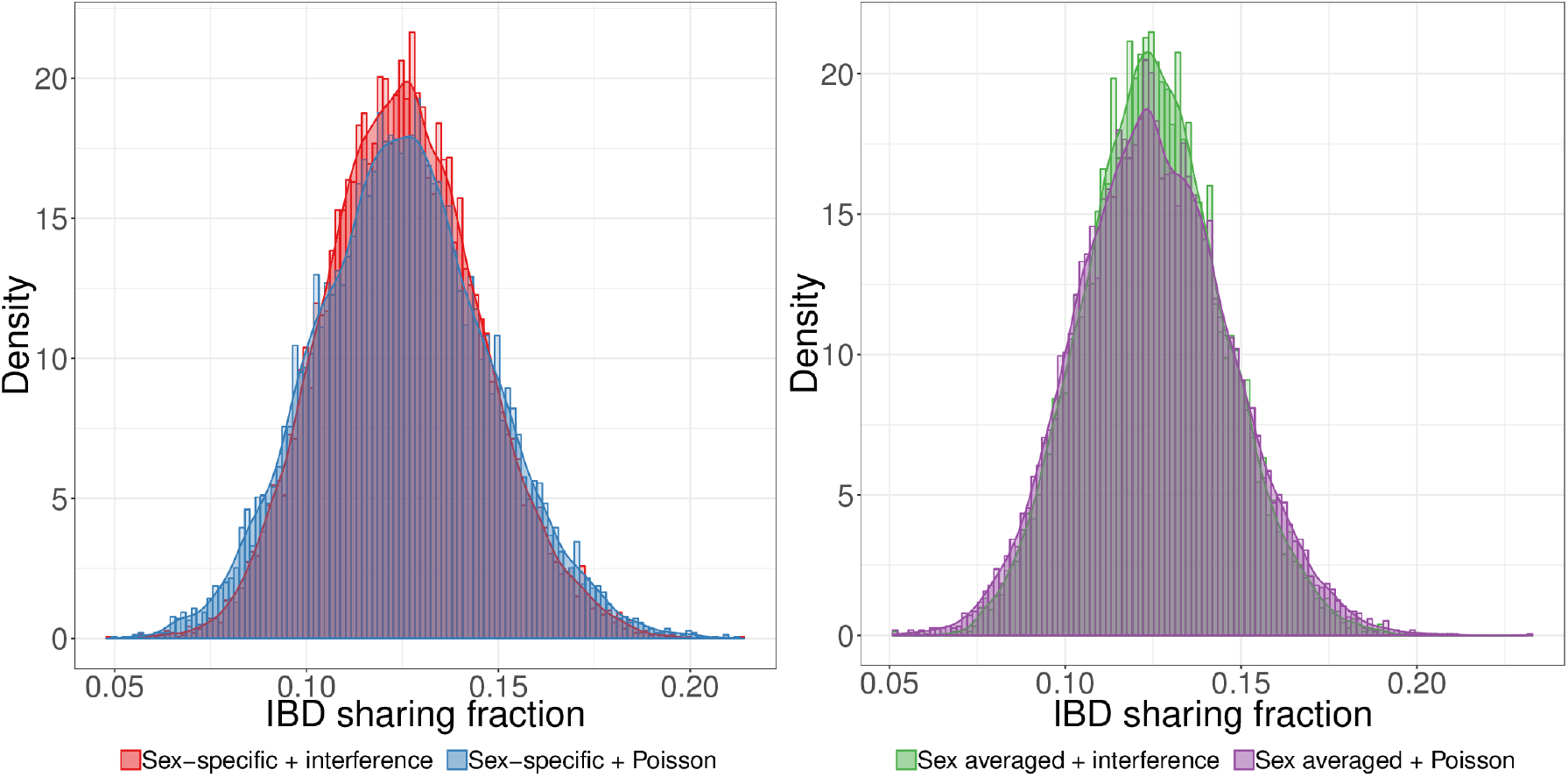
First cousins simulated with crossover interference have a distribution of IBD sharing proportion more concentrated near the mean than those simulated using a Poisson model. Interference decreases the variance in IBD sharing both when using sex-specific (left) and sex averaged (right) genetic maps.

### Simulations including sex-specific maps and interference best fit data from real relatives

Given the differences in the distribution of IBD proportions observed by varying the combination of map type and crossover interference among simulated relatives, we sought to understand which scenario best matches real human data. We first examined IBD sharing between pairs of full siblings in the SAMAFS and Hemani20k data, where use of family-based phasing enables IBD detection with high precision and recall. The mean IBD proportion in the SAMAFS data is 0.500, in line with the expectation, and the mean in the Hemani20k pairs is 0.502 (S1, S3 Fig). Overall, the SS+intf model produces the best fit to the standard deviations from real data, being the only model within one standard error (0.36 units) of the SAMAFS estimate, and 1.01 standard errors from the Hemani20k value (Fig 1, S4, S5). This contrasts with the traditional SA+Poiss model, which is 2.9 standard errors from SAMAFS, and 9.8 from Hemani20k. The SA+intf and SS+Poiss models are also discrepant, with both more than 8.3 standard errors from SAMAFS, and 5.4 standard errors from Hemani20k.

The mean IBD2 sharing rate in the SAMAFS full siblings is 0.250, as expected, and the corresponding value in Hemani20k is 0.251 (S1 Fig). The standard deviation of IBD2 sharing under the SS+intf model is 1.6 standard errors from that of SAMAFS, and only 0.55 standard errors from the Hemani20k value. These deviations are the smallest of all the models we considered (Fig 1). The traditional SA+Poiss model is the next closest to SAMAFS at a distance of 2.9 standard errors, but deviates meaningfully from Hemani20k at 9.8 standard errors away. The SA+intf IBD2 standard deviation is 7.2 and 3.5 standard errors from the SAMAFS and Hemani20k quantities, respectively, and, as in the full IBD proportion, SS+Poiss deviates the most from the real data, being 7.8 standard errors from SAMAFS, and 15.1 from Hemani20k.

Turning to relationships more distant than full siblings, we focus on the IBD sharing rates between the first and third quartiles, and we compare these to mean-shifted SAMAFS values. Additionally, we analyzed a subset of SAMAFS samples where all first degree relatives that connect them have available data that confirms their first degree status (S6A Fig, Methods). This subset should be free of any mislabeled relatives, and we refer to it as *SAMAFS-validated*.

As in the full sibling analyses, use of SS genetic maps and crossover interference modeling provides a good fit to the real data across all these more distant relationship types. In first cousins, the 25th and 75th percentile IBD proportions under the SS+intf model are 0.111 and 0.138—the same as in the SAMAFS-validated data (Fig 1)—while the corresponding percentiles under SA+Poiss are 0.110 and 0.140, with the latter value 3.7 standard errors from SAMAFS-validated. For first cousins once removed and second cousins, the SS+intf 25th and 75th percentile values are within one standard error of SAMAFS-validated in all cases, while the three other models deviate by more than one standard error for at least one of the two quartiles in both relationships.

As another line evidence that IBD sharing in SS+intf most closely mirrors that of real data, we inferred degrees of relatedness for the simulated and SAMAFS relatives. This inference maps the kinship coefficient of each pair to a degree of relatedness using the same kinship ranges as in KING [37] (Methods). S7 Fig plots the percentage of samples inferred as their true degree of relatedness in the SAMAFS and simulated pairs. For all four relationship types, the model with percentages nearest to that of SAMAFS-validated is SS+intf. In fact, SS+intf is within one standard error of the SAMAFS-validated percentage for all four relationship types, whereas SA+Poiss and SS+Poiss are >3.1 standard errors from SAMAFS-validated for all but full siblings. The SA+intf model is less than one standard error away from SAMAFS-validated for all but first cousins once removed, where it deviates by 2.7 standard errors.

### Rates of sharing at least one IBD segment among distant relatives

Random assortment during meiosis commonly leads to a loss of IBD segments such that distant relatives may not share any IBD regions with each other despite having a genealogical relationship. Given the fit of the crossover model that incorporates SS maps and interference, we set out to examine the distribution of the number of IBD segments shared among full and half-siblings and first through sixth cousins. For close relatives, including full and half-siblings, and first and second cousins, all simulated pairs share at least one IBD segment with each other regardless of the crossover model. However, some proportion of third through sixth cousins share no IBD segments of any size (Fig 3). Specifically, in the SS+intf simulation, 1.5% of third cousins share no IBD regions, and this percentage increases to 27.3%, 67.4%, and 88.9% of fourth, fifth, and sixth cousins, respectively. For the 1,112 (of 10,000) sixth cousins that do share IBD segments, the average total length is 7.6 centiMorgans (cM). Unsurprisingly, most sixth cousin pairs retain only one IBD segment with very few (107/1112) pairs sharing more than one segment (Fig 3). The total IBD length varies substantially among sixth cousins, with the top 25% of pairs that have IBD regions sharing a total of at least 10.2 cM and a maximum of 53.4 cM. Thus sixth cousins with rare extremes of IBD sharing have total shared lengths more typical of third and fourth cousins.

**Figure 3:**
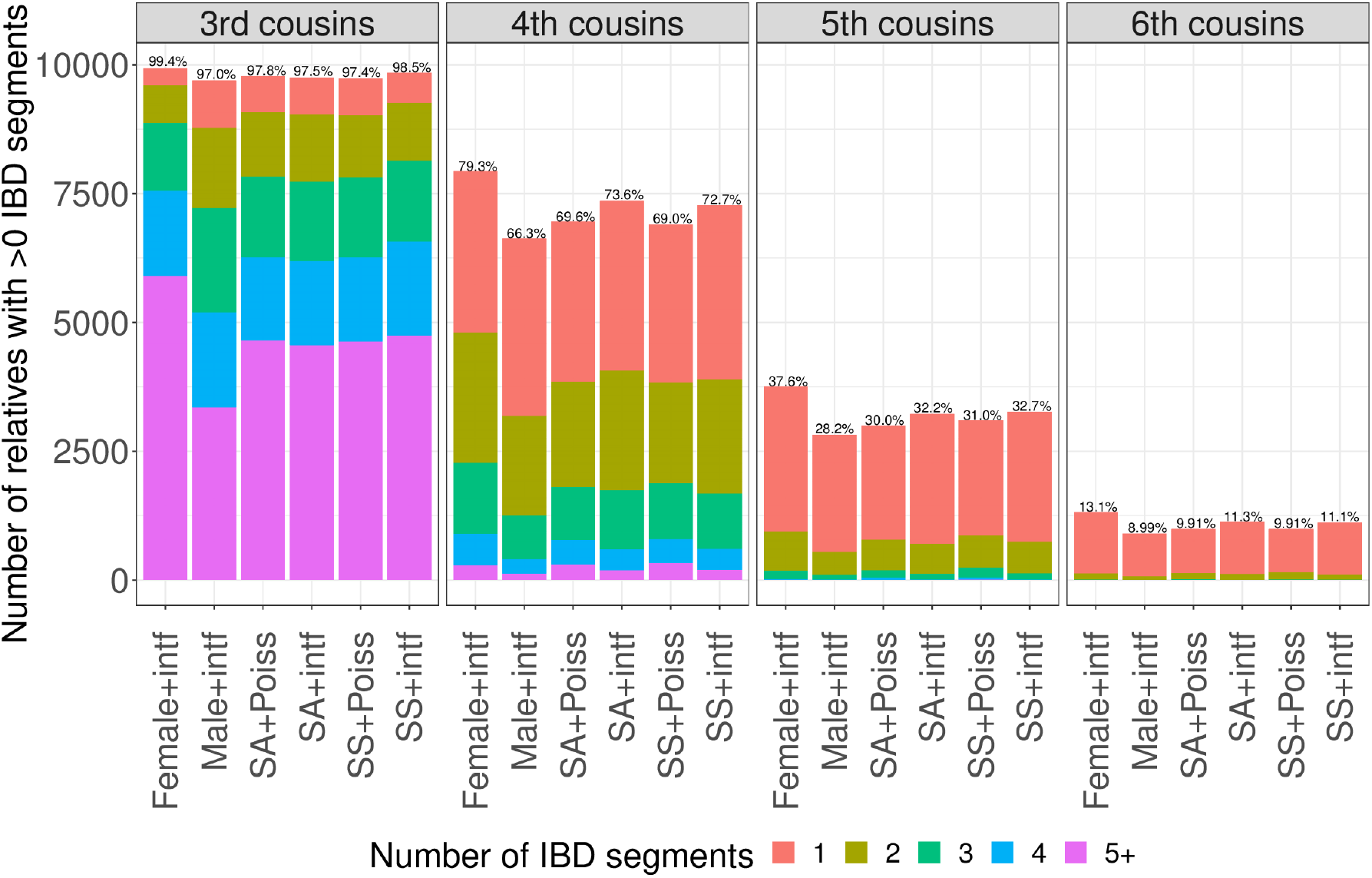
Number of IBD segments shared between simulated third through sixth cousins under various modeling scenarios. More distant relatives have reduced rates of sharing one or more IBD regions. Percentages above each bar indicate the fraction of simulated relatives (of 10,000 for each scenario) that have at least one segment shared. Female+intf are from simulations using sex-specific maps and interference but where the pairs are related through only female non-founders, with a male and female couple as founder common ancestors (S6B Fig). Male+intf pairs are the same as Female+intf but with the non-founders being only male instead of female.

As already noted, crossover interference leads to a more concentrated distribution of IBD sharing rates (e.g., Fig 2). Interference also leads to a slightly larger fraction of distant relatives that share IBD segments. For example, 32.7% of fifth cousins share one or more IBD segments under the SS+intf model compared to only 30.0% under SA+Poiss.

### Sex-specific maps dramatically impact the number of IBD segments relatives share

While SS maps have a smaller effect than interference on the variance in IBD sharing proportion between two relatives, they do impact the number of segments relatives share. Specifically, females produce an average of 1.57 times more autosomal crossover events per meiosis than males [16]. With such differences, females should transmit a larger number of IBD segments that are on average smaller compared to transmissions from males. This is because, without a crossover event, the probability of transmitting an IBD segment is 50%. On the other hand, when a newly generated crossover occurs within an IBD region, transmission of some portion of the IBD region (on one side or the other of the crossover) is guaranteed.

To more fully investigate the impact of SS genetic maps, we used the SS+intf model to simulate third through sixth cousins where the non-founder ancestors through whom they are related are either all female or all male (with the shared founder grandparents being a male and female couple; S6B Fig). When related primarily through females, third through sixth cousins are much more likely to share some amount of IBD than those related primarily through males. The differences are quite extreme with respectively 2.5%, 19.6%, 33.1%, and 46.2% more (in relative terms) third, fourth, fifth, and sixth female-lineage cousin pairs sharing at least one IBD region compared to the analogous male-lineage cousins (Fig 3). Consistent with intuition, the IBD regions in female-descent cousins are smaller on average than those in male-descent cousins. For example, female-lineage fifth cousins with IBD regions share an average of 1.3 segments with a mean total length of 9.0 cM compared to the male-lineage averages of 1.2 segments and total length 11.9 cM.

These differences in male and female maps impact IBD sharing between close relatives as well, with especially noticeable effects in half-siblings. In particular, maternal half-siblings share on average 1.4 times as many IBD segments as paternal half-siblings (mean segment numbers 51.9 and 37.1, respectively). The effect is substantial enough to produce a bimodal distribution, with little overlap between the two types of half-siblings (Fig 4; S8A). Although less distinct than segment counts from simulations, the SAMAFS half-siblings also have a bimodal distribution that corresponds with the sex of the common parent (S8B Fig; Methods). Notably, the mean segment count in simulated paternal half-siblings is less than that of first cousins with randomly assigned parent sex (who share a mean of 39.0 segments; Fig 4). However, the segments paternal half-siblings share are more than twice as long as those of first cousins, with an average length of 45.1 cM compared to 21.5 cM in first cousins.

**Figure 4:**
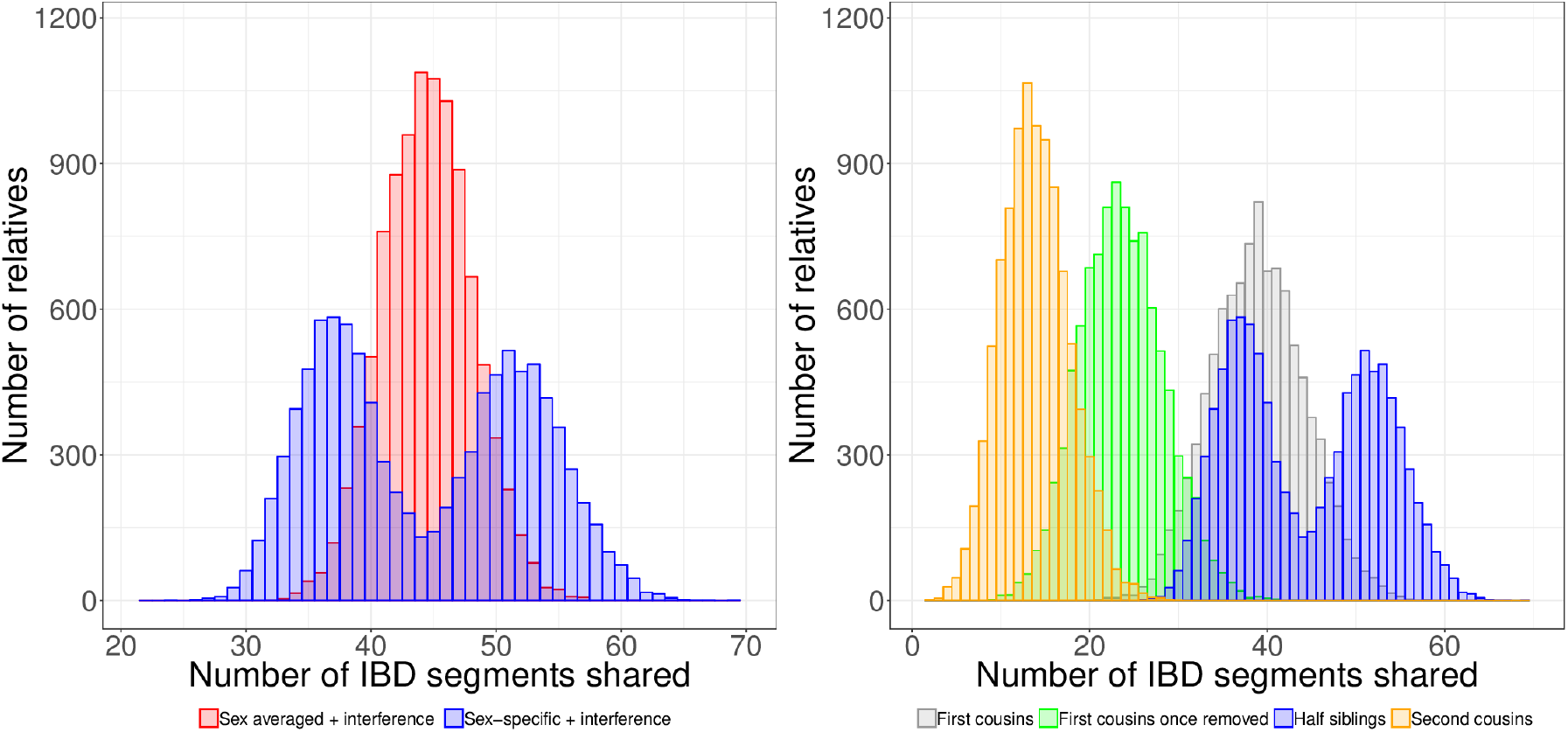
Sex-specific maps impact the number of segments half-siblings share. Number of IBD segments half-siblings share when simulated with sex averaged maps compared to sex-specific maps have very different shapes, with sex-specific maps producing a bimodal distribution (left). Half-sibling segment counts in the context of other relative types also simulated under sex-specific maps (right). The lower mode of half-sibling segment counts—which corresponds to IBD sharing between paternal half-siblings (S8A Figure)—is below that of first cousins. The distributions are based on 10,000 pairs simulated under interference for all relationship types.

### Estimates of time since admixture vary more under sex-specific maps

We sought to characterize the impact of crossover models on estimates of the time since admixture using single admixed samples. For this purpose, we treated IBD segments shared by a given individual and one of their (*T* + 1)^th^ generation ancestors as local ancestry tracts. We then used all the segments the pair shares to estimate the time since admixture by fitting an exponential rate. Another way to view this is as an estimate of how many meioses separate an individual from a given direct ancestor. Under both perspectives, the first meiosis is unobserved because, in the context of admixture, an unadmixed ancestor transmits a chromosome that is entirely composed of one ancestry in the first meiosis (the crossovers do not yield ancestry switches). For estimating the number of meioses to an ancestor, the crossovers in the first meiosis affect which haplotype the ancestor transmits, but population-based IBD methods typically infer segments that span both haplotypes and do not identify switches between haplotypes. (In principle, well-phased data could enable localization of IBD segments to specific haplotypes of an ancestor, yet phasing of this quality is only feasible using data from families.)

Fig 5 plots estimates of *T* based on exponential rate fits in 10,000 individual-ancestor pairs. Here, crossover interference has a limited effect on the distribution, leading to a reduction in standard deviation compared to the Poisson model of 5.9% (averaged over *T* =1, 2, 3, 5). By contrast, the variance under SS maps is much higher than under the SA map for *T* =1 (a grandparent-grandchild pair) and *T* = 2 (great grandparent-grandchild pair), with standard deviations 1.37 and 1.20 times larger under the SS maps, respectively. For *T* = 3 and *T* = 5, the standard deviations under the SS maps remain larger than under the SA map (1.08 and 1.34 larger, respectively), but the effects on the bulk of the distribution are limited. Specifically, the interquartile ranges for *T* = 3 and *T* = 5 meioses are only 1.12 and 1.02 times larger, respectively, compared to the SA map. The differences between the SS and SA models are highest for small numbers of meioses because the probability of all meioses being in only one sex is highest for smaller numbers of meioses. As the number of meioses grows, a greater fraction of the samples will have closer to equal numbers of male and female meioses, and so the sharing patterns will be more similar to those that arise from an SA map.

**Figure 5:**
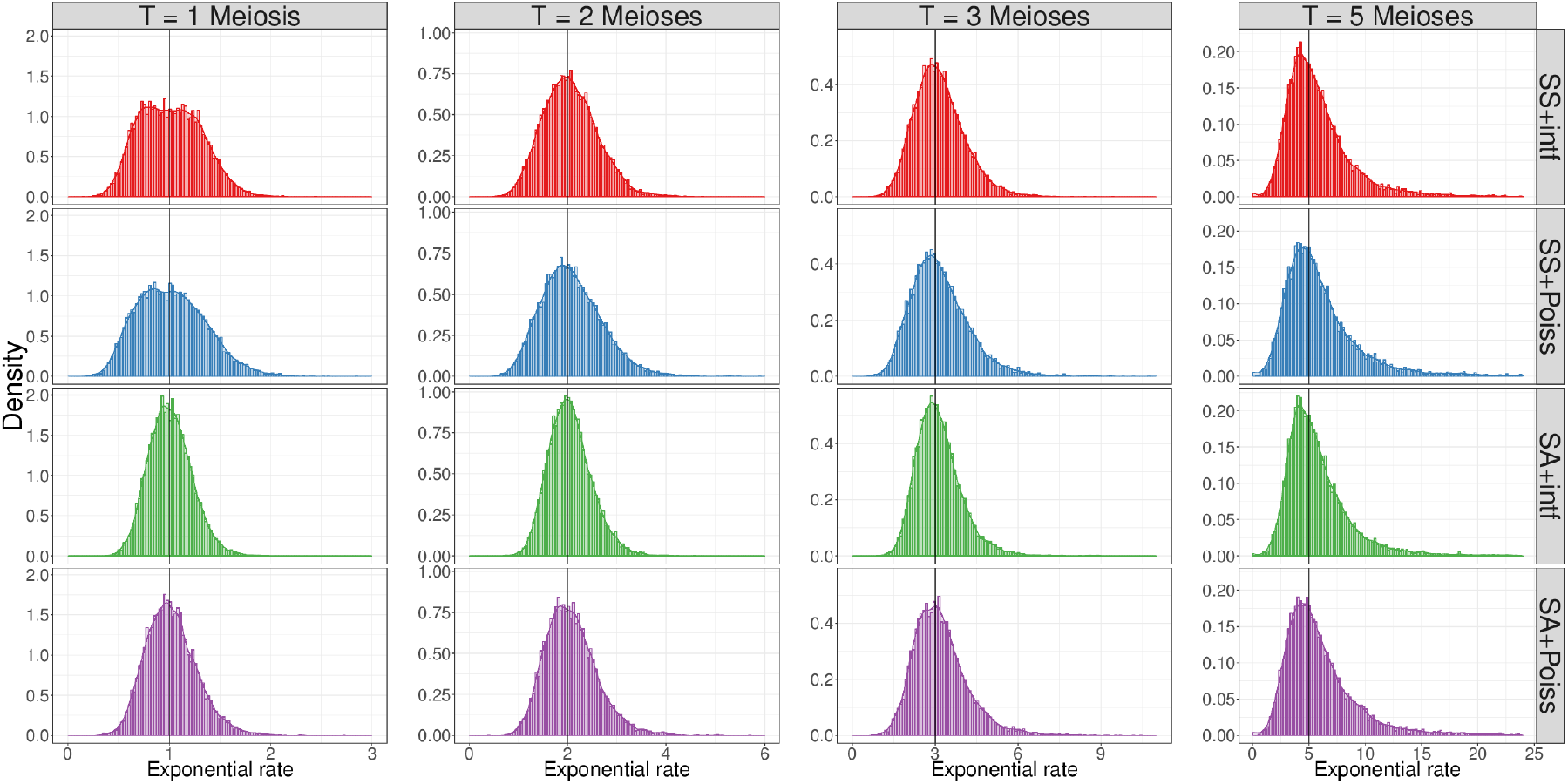
Distribution of estimated time since admixture based on one admixed sample. Histograms show estimated rates for 10,000 individuals under each simulation scenario. Estimates are rate fits of an exponential distribution accounting for finite chromosomes (Methods). Horizontal lines indicate the true *T*.

The above analysis focuses on only one (logically) admixed sample. When considering segments from all 10.0 pairs, the shapes of the segment length distributions vary—being especially altered by interference for small *T*—yet the exponential rates are effectively identical across the simulation models (S9 Fig). These identical rates reflect the fact that the crossover rates (map lengths) are the same across all models when averaged between the sexes. Thus, given sufficient data, time since admixture estimates appear unaffected by either interference or SS maps.

### The effects of interference and sex-specific maps on IBD segment lengths

To gain insight into the effect of crossover interference and SS maps on IBD segment lengths, we analytically obtained the distribution of these lengths under the SA+intf and SS+Poiss models.

In the Housworth-Stahl two-pathway model [14], the proportion of crossovers that escape interference (are “unregulated”, or distributed according to a Poisson process) is denoted *p*. The remaining (“regulated”) crossovers are independently generated by first drawing the positions of chiasmata as a stationary renewal process [38] along the chromosome, with gamma distributed inter-chiasma distances (in Morgans) with shape *ν* and rate 2*ν*(1 − *p*) (Methods). Each chiasma becomes a crossover in the gamete being modeled with probability 1/2. Here we assume that two chromosomes are taken from individuals with a common ancestor *T* generations ago, or separated by 2*T* meioses.

In Methods, we show that the density of *x*, the length (in Morgans) of IBD segments subject to interference and under an SA map, is

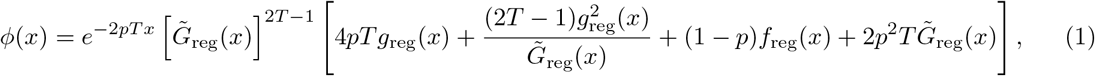

where

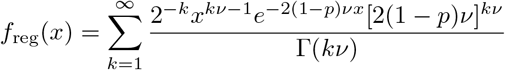

is the probability density of the distance between regulated crossovers,

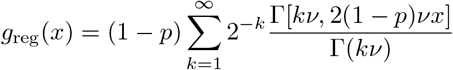

is the probability density of the distance between a random site and the next regulated crossover, and

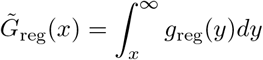

is one minus the cumulative distribution of *g*_reg_(*x*).

The expressions above are valid for infinitely-long chromosomes. We further show in Methods how to modify Eq. (1) for the case of a finite chromosome (Eq. (16)).

To confirm these results, we used Ped-sim to simulate IBD sharing under the SA+intf model for chromosome 1. The simulated distribution of the IBD segment lengths is shown in Fig 6 for half-cousins with a common ancestor *T* = 1, 2,4, 6 generations ago (where *T* = 1 corresponds to half-siblings), and shows agreement with the theory (Eq. (1)). The plot also depicts the expected distribution under the Poisson model, and demonstrates that the effect of interference can be substantial and is noticeable up to *T* ≲ 4. However, by *T* = 6 the Poisson process is already an excellent approximation to SA+intf.

**Figure 6:**
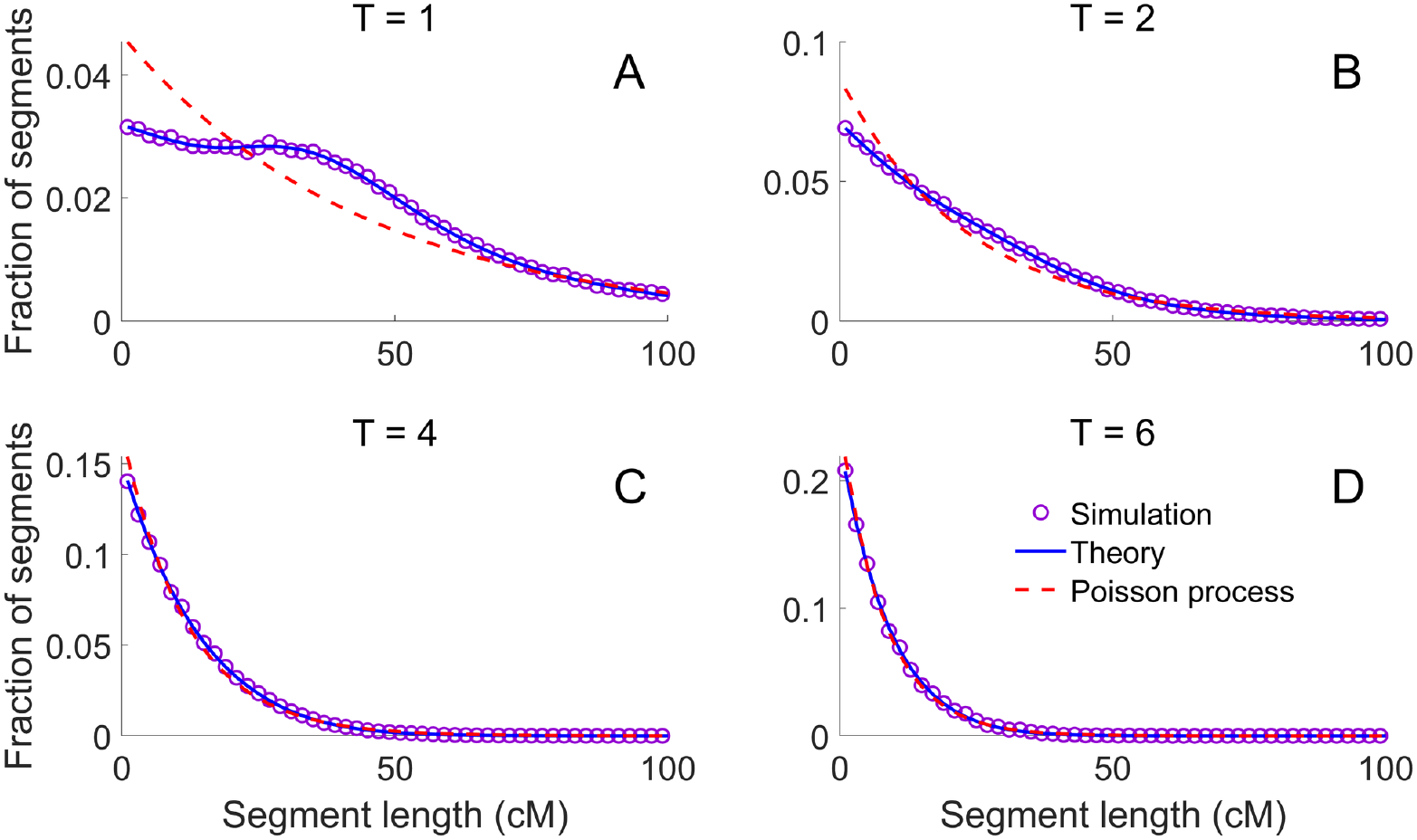
The effect of crossover interference on IBD segment lengths. We used Ped-sim to simulate half-cousins with a common ancestor *T* = 1, 2,4, 6 generations ago under the SA+intf model, extracting IBD segment lengths for chromosome 1 (panels A-D, respectively). Each panel shows the simulated distribution of IBD segment lengths (over 10^5^ pairs for *T* = 1, 2 and 10^6^ pairs otherwise; purple circles), the theory from Eq. (1) (blue lines; includes the finite-chromosome correction of Eq. (16)), and the expectation based on a Poisson process (red dashed lines; Eq. (17)).

Next we considered the effect of SS maps, this time assuming Poisson crossover placement. For concreteness, consider (*T* − 1)^th^-full cousins, which are separated by 2*T* meioses as above. Each IBD segment descends from one of the founder parents (a female or a male with equal probability), who transmits it via two meioses. The remaining 2*T* − 2 meioses can be either male or female with equal probability. The number of female transmissions is thus *n_f_* = *n_f,i_* + 2*n_f,a_*, where *n_f,i_* is the number of female meioses that relate the full cousins ignoring the common ancestors, and is binomial with parameters (2*T* − 2,1/2); and *n_f,a_* indicates whether the common ancestor who transmitted the segment is female, and is Bernoulli with parameter 1/2. The number of male transmissions is *n_m_* = 2*T* − *n_f_*. In fact, the same expressions hold for half-cousins, if the sex of the founder parent is random.

For each SNP *i*, denote by λ_*m*_(*i*) the male crossover rate (in Morgans per base-pair [bp]) between SNPs *i* and *i* + 1, and define λ_*f*_(*i*) similarly. We assume the rate is constant between SNPs (and zero before the first SNP). Given *n_f_* and *n_m_* female and male meioses, respectively, the total crossover rate between SNPs *i* and *i* + 1 for the two relatives is λ(*i*) = λ_*f*_(*i*)*n_f_* + λ_*m*_(*i*)*n_m_*. Thus, placement of crossovers is still based on a Poisson process, but because the per bp rates in males and females differ by position, the rate is inhomogeneous along the genome. (Note that the male and female maps themselves are also inhomogeneous with respect to physical positions. We focus here on physical positions because the effects of the process ultimately occur at a physical position, and those physical positions are common to both the male and female maps.)

To obtain the distribution of inter-crossover distances (again in physical bp) for a fixed number of male and female transmissions, we use a result by Yakovlev *et al*. [39] for the distribution of inter-event times in an inhomogeneous Poisson process. Denote λ(*x*) as the implied crossover rate at a physical coordinate *x* (as implied by the λ(*i*)’s above), and define 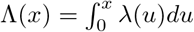. Then we have

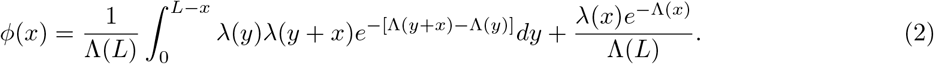

Here *ϕ*(*x*) describes the density of all inter-crossover distances, not including the one censored by the chromosome end, with the number of male and female transmissions assumed given. To obtain the density without conditioning on male and female meiosis counts, we sum over all *n_f_* = 0,…,2*T*, each weighted by its probability. In relatives, not all inter-crossover blocks will become IBD segments, but rather only those whose line of descent is from the same common ancestor in both relatives. However, since IBD segments are a random subset of all blocks, the IBD segment length distribution is expected to be similar to that of the inter-crossover distances. This is confirmed in S10 Fig, where we plot the distribution of simulated IBD segment lengths under SS+Poiss for half-cousins separated by *T* = 1, 2,4, 6 generations. As opposed to the observations from crossover interference, the distribution of segment lengths under SS maps is not substantially different from that obtained by Eq. (2) with SA maps.

### Ped-sim comparison to IBDsim

The functionality in Ped-sim is available in other packages, though with some limitations. To our knowledge, the most similar method is IBDsim [40], an R package that uses the *χ*^2^ interference model with fixed parameters. We used Ped-sim (under the SS+intf model) and IBDsim to simulate 10,000 full sibling pairs and 10,000 second cousin pairs (Methods). Ped-sim simulated the full siblings and second cousins in 8.1 and 8.7 seconds, respectively, while IBDsim required 371 and 608 seconds, respectively (corresponding to ~45-fold and ~68-fold speedups). Memory requirements are low for both methods, with Ped-sim and IBDsim using, respectively .62 Gb and 2.0 Gb to simulate the second cousins.

Neither of the above analyses produced genotype data (and IBDsim does not provide this functionality), but only generated IBD segments from replicate pedigrees. To produce genotype data, Ped-sim compute times are on the scale of dozens of minutes for several thousand samples. For example, simulating genotype data for 4,450 pairs of full siblings at 462,828 markers took 69.9 minutes and used .14 Gb of RAM.

## Discussion

Modeling relatedness among individuals is more challenging than is typically appreciated due to the complexities of meiotic biology. Variable crossover rates between the sexes and the phenomenon of crossover interference both affect the quantity and size of IBD segments that individuals share. Our analyses demonstrate that use of sex-specific maps and inclusion of crossover interference provides the best fit to the standard deviation of IBD sharing rates in real human data from full siblings. Likewise, the 25th and 75th percentiles of IBD proportion from real first through second cousins are best fit by jointly modeling sex-specific maps and interference.

In modeling both the IBD sharing proportion between relatives and the lengths of their IBD segments, crossover interference has a much stronger influence than varying sex-specific versus sex averaged maps. However, sex-specific maps have a sizable impact on the number of IBD segments that both close (especially half-siblings) and distant relatives share. Therefore, both crossover interference and sex-specific maps play critical roles in the accurate representation of meiotic transmissions. Even so, the crossover model that is most discrepant with real IBD sharing fractions adopts sex-specific maps but Poisson inter-crossover distances. Thus, although closer to the true meiotic process in terms of the number of crossover events in men and women, the strong impact of interference in reducing the variance of IBD sharing is important to include when simulating relatives under a sex-specific map.

Given the effects on IBD sharing of the features we consider here, it is necessary to revisit the probability that a pair of relatives share any IBD with each other. A classic, influential treatment of this problem used an analytical approach based on Markov models and considered sex averaged maps while ignoring interference [18]. That study indicated that 10.1% of sixth cousins share IBD regions, which is close to the 9.9% we obtain using a more up-to-date sex averaged genetic map. Still, with both sex-specific maps and interference modeling, we find that 11.1% of simulated sixth cousins have non-zero IBD sharing. This factor rises to 13.1% when the sixth cousins are related primarily through females, and drops to 9.0% when they are related primarily through males.

Overall, these findings are practically important in the context of relatedness inference (S7 Fig), estimates of the time since admixture (Fig 5, S9), and linkage analysis. As a primary basis of genetic genealogy, and standard practice in genetic testing companies, relatedness inference using data from consumer genetic testing kits is fueling a revolution in genealogical research. Analyses of the simulated relatives demonstrate that individuals are classified to their true degree of relatedness more often when simulated under interference than under the Poisson model (Figure S7). For example, 4.3% more second cousins simulated under the more refined model have kinship coefficients that map to their true degree of relatedness compared to those simulated with the standard sex averaged, Poisson model. Even so, for many applications, meiosis counts ≳ 12 (or *T* ≳ 6 in Fig 6, S10) between individuals are reasonable to analyze using the traditional model. That is, while crossover models affect IBD sharing in even moderately distant relatives such as fifth or sixth cousins, population-based coalescent simulations and inference techniques are generally reasonable to perform with standard approximations. Still, fine grained analyses of relatives as far distant as sixth cousins (or more distant) will benefit from the more precise models analyzed here. One such application is linkage analysis, where, given a known pedigree structure that includes the sexes of the individuals that relate a collection of genotyped samples, one can construct a more precise null model of IBD sharing using these models of crossovers.

Going forward, efforts to better understand the dynamics of crossovers, including observed “gamete effects” wherein crossover counts in a given gamete are correlated across chromosomes [41], and incorporating genetic variants that effect crossover rates [41] could yield models with even greater precision than those we considered. Nevertheless, the effects of crossover interference and sex-specific maps merit consideration in models of close relatives and alter IBD distributions in even moderately distant relatives.

## Methods

We analyzed data from a combination of simulated and real relatives, the latter from the SAMAFS and 20,240 full sibling pairs from Hemani *et al*. [34]. The IBD sharing statistic we focus on primarily is the proportion of their genome two relatives share IBD, calculated as a fraction of the diploid genome. For a given pair, this proportion is (*k*^(2)^ + *k*^(1)^/2), where *k*^(2)^ and *k*^(1)^ are the fraction of positions (in genetic map units) the pair shares IBD2 and IBD1, respectively. To perform degree of relatedness classification (S7 Fig), we used kinship coefficients calculated for a given pair as (*k*^(2)^/2+*k*^(1)^/4)—i.e., 1/2 the IBD proportion—and mapped these coefficients to degrees according to the ranges KING uses [37].

### The Ped-sim algorithm

Ped-sim simulates relatives by tracking haplotypes—initially ignoring genetic data—as a sequence of segments that span a chromosome. Each segment consists of a numerical identifier denoting the founder haplotype it descends from and a segment end point. (The start position is implicitly either the beginning of the chromosome or the site following the end of the previous segment.) All founders have two haplotypes with only one chromosome-length segment, each with a unique identifier.

To begin, Ped-sim reads a file that defines the pedigree structure(s) it is to simulate and, for each such structure, generates haplotype segments for the founders in the first generation. For subsequent generations, it generates haplotypes for any founders in that generation and forms haplotypes for non-founders from the parents’ haplotypes under a meiosis model. This model works on the two haplotypes belonging to a given parent by first randomly selecting which of these begins the offspring haplotype, each having 1/2 probability of being selected. Next, Ped-sim samples the location of the crossover events, either using a model of crossover interference or a Poisson model. It then produces the offspring haplotype by copying the segments that comprise the parent’s initial haplotype up to the position of the first crossover, and introduces a break point in the copied segment at that crossover position. Following this, it switches to copying segments from the parent’s other haplotype, and it continues to alternate copied-from haplotypes at each crossover in this manner until the end of the chromosome.

Details of the Housworth-Stahl crossover interference model are below, and we discuss parameter choices in the next subsection.

Under the Poisson crossover model, the distance from the start of the chromosome to the first crossover, and from one crossover to the next are each exponentially distributed with rate equal to 1 crossover/Morgan. This rate arises naturally from the definition of a Morgan as the distance within which an average of one crossover occurs per generation. The model sequentially samples crossovers and terminates after sampling a crossover beyond the end of the chromosome.

Both models produce crossover positions in genetic units (i.e., Morgans), and Ped-sim determines their physical location using a genetic map, storing the segment end points as physical bp positions. When using sex-specific maps, it locates the physical positions using the map corresponding to the sex of the parent. If a crossover falls between two defined map positions, Ped-sim uses linear interpolation to determine the physical location.

By default, Ped-sim randomly assigns the sexes of parents, and can generate any number of pedigrees with a given structure (with parent sexes assigned independently in each). Ped-sim can also generate data in which all reproducing non-founders have the same sex (male or female), leading to descendants that are related to each other through nearly all male or all female relatives. In such cases, the common ancestors of those descendants will generally be a married couple (S6B Fig), male and female, although it is possible to simulate only a single common ancestor or for individuals to be related through more than one lineage (e.g., double first cousins).

When given genetic data in the form of input haplotypes, Ped-sim randomly assigns the data from one input sample to each founder. It then copies alleles from these assigned founder haplotypes to descendants using the segment numerical identifiers and end points. The algorithm can also introduce genotyping errors and missing data using user-specified rates, with a uniform probability of these events at all positions.

Another way to run Ped-sim is without haplotype data—instead using only IBD segments detected using the internally tracked haplotype segments. This is the way we ran Ped-sim for the analyses we describe here, using version 1.0.1 of the tool. The IBD segments Ped-sim generates consist of physical start and end positions in both physical (bp) and genetic units (cM). To be able to analyze statistics of IBD sharing in terms of genetic distances, Ped-sim reports the sex averaged genetic start and end positions of each segment (the midpoint of the sex-specific coordinates), even when using sex-specific maps.

Ped-sim is available from https://github.com/williamslab/ped-sim, with several example pedigree definition (`def`) files and the interference parameters we used [15] included in the repository. The documentation in the repository includes links and Bash code for downloading and generating a file with the sex-specific map we used [16].

### Genetic maps and crossover interference parameters

We used genetic maps produced using crossovers from over 100,000 meioses [16]. These maps include those for both males and females and a sex averaged map that all span the same physical range. All simulations include the 22 autosomes but no sex chromosomes.

To simulate using the Housworth-Stahl crossover interference model, we leveraged female and male interference parameters *ν_i_* and *p_i_* for *i* ∈ {*f, m*}, respectively, that were inferred from over 18,000 meioses [15]. We calculated the sex averaged parameters *ν_a_* and *p_a_* as follows. The *p* parameter gives the fraction of events that escape interference, and we set *p_a_* = (*p_f_* + *p_m_*)/2. In this model, distances between chiasmata subject to interference are gamma distributed with shape and rate parameter values *ν* and 2(1 − *p*)*ν*, respectively [13, 14]. A simple average of the male and female *ν* parameters does not produce a distribution with summary statistics at the midpoint between the two sexes. All values of *ν* lead to distributions with the same expected value of 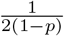 because the expectation of a gamma distribution is the shape divided by the rate parameter. We therefore calculated *ν_a_* such that the variance of the sex averaged distribution is the mean of the variances of the male and female models, while assuming all *p* parameters are the same between the models (since we separately estimate *p_a_*). This gives 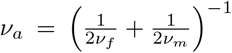. Note that the mean distance of 1/2 Morgans between events (accounting for both the regulated crossovers and those that escape interference) is half the distance expected per chromatid. This is because the model is for events in a tetrad (all four products of meiosis). As noted earlier, to obtain the crossovers falling on the gamete being generated, the model randomly selects events with probability 1/2 for inclusion.

### IBD detection in the SAMAFS

We used two different methods for inferring IBD fractions in the SAMAFS data: one applied to nuclear families and which we used to analyze IBD sharing between full siblings, and the other for analyzing IBD rates in first cousins, first cousins once removed, and second cousins. Quality control filtering of the SAMAFS data is the same as that described previously [42, 43]. In brief, we used biallelic SNPs typed on the Illumina Human660W, Human1M, Human1M-Duo, or both the HumanHap500 and the HumanExon510S arrays, and required the SNP probe sequences to map to a single location in the human GRCh37 build. Next, we excluded individuals and SNPs with excessive missing data (>10% and >2%, respectively) and removed duplicate SNPs. Additional SNP filters utilized information from auxiliary resources including dbSNP and the reported “accessible genome” from the 1000 Genomes Project, among others [42]. This yielded data for 2,485 samples typed at 521,184 SNPs. We further omitted 1,514 first cousin, first cousin once removed, and second cousin relative pairs that had evidence of being related through more than one lineage [43].

Family-based phasing implicitly infers IBD regions, and in the presence of data for a complete nuclear family, this inference has both high precision and recall. For this analysis, we utilized HAPI [35] version 1.87.6b, a method that performs efficient minimum recombinant phasing for nuclear families. This form of phasing is the same as that of the Lander-Green algorithm [44] when the probability of crossover between informative markers is identical at each position. To ensure reliable results, we performed this inference on 116 nuclear families for which data from both parents and three or more children were available, and we excluded one member of likely monozygotic twin pairs that had IBD2 rates >0.95 (three pairs).

To infer IBD regions, we parsed the inheritance vector output from HAPI to locate IBD1 and IBD2 segments, assigning genetic positions to the start and end of each such region using the same sex averaged map we used for the simulated data [16]. (The genetic map is undefined for 102 SNPs and we omitted these positions from analysis.) The exact boundaries of crossover positions are uncertain in real data due to the fact that not all sites are genotyped and homozygous positions are uninformative. We therefore estimated the start and end positions as the midpoint in genetic units between two informative sites that descend from distinct parental haplotypes and therefore bound the region in which a crossover broke an IBD segment. To ensure the IBD intervals are real and not due to genotyping error, we also merged short regions (including non-IBD intervals) comprised of ten or fewer informative SNPs with the adjacent segments so that they cover the interval. We assign these to have the same IBD type as the preceding segment (typically the two flanking segments are of the same type). On average each pair had 4.7 merged regions across all autosomes (standard deviation 10.8). We removed pairs that had more than 26 merged regions (i.e., more than two standard deviations from the mean). This filter removed a total 16 of pairs, and 15 of these were from five families. We therefore removed these families from the analysis, leading to the further exclusion of 13 full sibling pairs. Finally, we removed two outlier full sibling pairs that had low IBD proportions of 0.358 and 0.371. This yielded 1,114 pairs of full siblings for analysis.

For non-sibling relatives, we leveraged IBD segments previously inferred [43] using Refined IBD [36] version 4.1. In total, the SAMAFS relatives consist of 5,384 pairs of first cousins, 6,342 first cousins once removed, and 2,584 second cousins. Here as well we converted physical positions of the IBD segments to sex averaged genetic positions using the same sex averaged map as in other analyses [16].

While the SAMAFS relationships are reliable [43], the statistics we consider are sensitive and can be meaningfully affected by small numbers of mislabeled pairs. We therefore sought to identify a *validated* set of relative pairs whose relationships are nearly certain to be correct. For validation, we required pairs to be descended from full siblings that are genotyped and inferred as first degree relatives by Refined IBD. We further required data for all parent-child relatives that fall between those full sibling ancestors and their descendants, and ensured that Refined IBD inferred the parent-child pairs as first degree relatives (S6A Fig). Subsetting the data in this way produces a validated set of 3,722 first cousins, 2,869 first cousins once removed, and 906 second cousins. Only one pair of the full sibling ancestors were not inferred as first degree relatives by Refined IBD, while Refined IBD inferred all the relevant parent-child pairs as first degree relatives.

We noted that the mean IBD rates for the real relatives are slightly elevated, potentially due to background relatedness or false positive IBD segments. For first cousins, the mean amount of IBD shared exceeds the theoretical expectation by 11.3 cM, and 19.5 cM in the validated set (0.17% and 0.29% above the expectation, respectively). For first cousins once removed, the observed means are greater than supported by theory by 0.17 cM (0.0026%), and 15.9 cM (0.24%) for the validated set. Finally, the excess for second cousins is 17.6 cM (0.26%), and 6.8 cM (0.10%) for the validated set. We subtracted off these mean excesses for each of these (non-sibling) relationship types in Fig 1 and S1 Fig and associated analyses; the unmodified statistics are in S1 Dataset. We did not mean-shift the kinship coefficients used to map the SAMAFS samples to degrees of relatedness (S7 Fig).

### IBD detection in SAMAFS half-siblings

To detect IBD segments shared between SAMAFS half-siblings (S8B Fig), we identified pairs listed as half-siblings in the SAMAFS pedigrees and contained in the 2,485 samples used for other analyses. Next, we retained pairs Refined IBD inferred as second degree relatives [43]. We then ran IBIS [45] to detect IBD segments using genetic positions from the SA map and parameter settings of: minimum segment length of 7 cM, window size of .01 cM, and an error threshold of .04. Finally, we merged adjacent segments shared between a given pair of individuals if the gap between the segments was ≤ 0.5 cM.

### Estimating time since admixture corrected for finite chromosomes

To estimate the time since admixture based on IBD segment lengths, we assumed the segments derive from a Poisson process—and thus segment lengths follow an exponential—and used the following maximum likelihood approach. The likelihood of a segment of length *x* is *Te*^−*Tx*^. However, segments bounded by the end of the chromosomes are “censored,” and we only know they are longer than *x*, and thus they have likelihood *e*^−*Tx*^. Assuming the segments are independent, the likelihood of all segments is 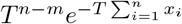, where there *n* segments of lengths *x*_1_,…,*x_n_, m* of which are censored by chromosome ends. Equating the derivative of the log-likelihood to zero, we obtain the maximum likelihood estimate 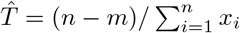.

### Deriving the distribution of IBD segment length under the Housworth-Stahl interference model Background

We assume a sex-averaged genetic map and that the chromosome is infinitely long (we will relax this assumption later). Under the Housworth-Stahl two-pathway model [14], a proportion *p* of crossovers escape regulation (referred to as “free” crossovers below), and are thus distributed along the chromosome as a Poisson process with rate *p* per Morgan. Regulated crossovers are generated independently of the unregulated crossovers by first drawing the positions of chiasmata as a stationary renewal process [38] along the chromosome, with gamma distributed inter-chiasma distances (in Morgans) with shape *ν* and rate 2*ν*(1 − *p*). Then, each chiasma becomes a crossover on the modeled gamete with probability 1/2, since it affects only one of the two sister chromatids. The latter process is called *thinning* and assumes no chromatid interference. The average inter-crossover distance is 1 Morgan, as per the definition of the Morgan unit.

In the regulated process, if *k* − 1 chiasmata are skipped between crossovers, the distance to the next crossover is distributed as gamma with shape *kν* and rate 2(1 − *p*)*ν*. After thinning, the distance between regulated crossovers is distributed as

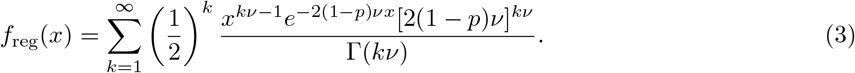

The free process has inter-crossover distances distributed as

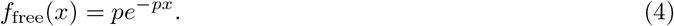

Below, we study the distance between crossovers across 2*T* meioses. As explained in Results, the IBD segment length distribution is expected to be similar to that of the inter-crossover distance distribution, as IBD segments are a random subset of the inter-crossover regions. To proceed, we will first compute the properties of the distance from a fixed point to the nearest downstream crossover (initially in one meiosis, then in multiple meioses), and then use these results to derive the distribution of inter-crossover distances across multiple meioses.

### The distance from a fixed point to a crossover in one meiosis

The process of placing chiasmata is assumed to be a stationary renewal process [14]. In one meiosis, the distance between a randomly selected site and the next crossover to the right (or left) due to the regulated process is distributed as [38]

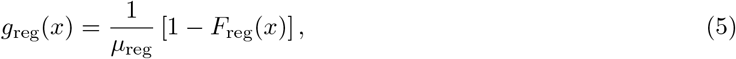

where *F*_reg_(*x*) is the CDF of *f*_reg_(*x*), i.e., 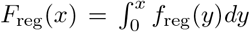, and *μ*_reg_ is the mean distance between regulated crossovers (i.e., 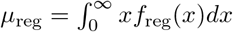). As *μ*_reg_ = 1/(1 − *p*), we have

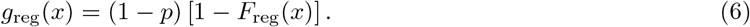

This can be written explicitly as

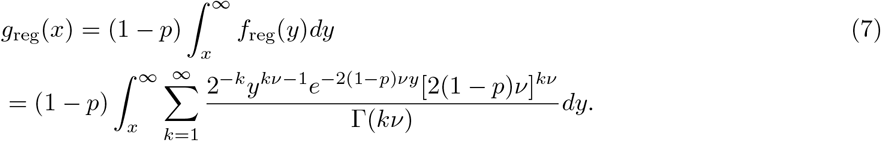

Changing variables, *z* = 2(1 − *p*)*νy*, we obtain

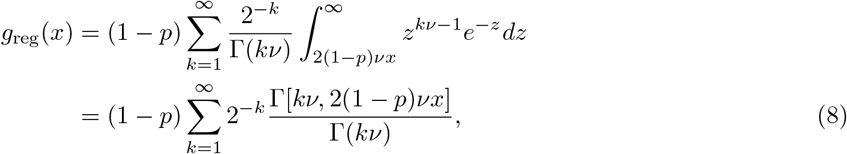

where Γ[*a, x*] is the upper incomplete gamma function. We denote the CDF of *g*_reg_(*x*) as *G*_reg_(*x*). For the free process, as it is memory-less,

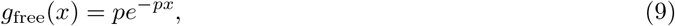

and the CDF of *g*_free_(*x*) is

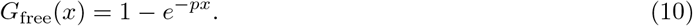

### The distance from a fixed point to a crossover across multiple meioses

The previous subsection described the distance to the next crossover of a given type (regulated/free) for a single meiosis. Given a focal site, the distance to the next crossover across 2*T* meioses and both processes is the minimum of the distance to 2*T* regulated processes and 2*T* free processes. We model all processes as independent, which is clearly true for meioses in different individuals, and holds for the regulated and free processes under the Housworth-Stahl model. Thus, the distance to the next crossover across 2*T* meioses is distributed as

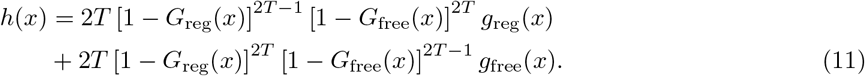

In the first term, all free crossovers and all but one of the regulated crossovers have distance larger than *x*, and one of the regulated crossovers has distance *x*. There are 2*T* possibilities to choose which regulated crossover has the minimal distance, and hence the initial factor of 2*T*. The second term is similar, with the minimal distance now coming from one of the free crossovers. Eq. (11) can be simplified based on *g*_free_(*x*) and *G*_free_(*x*) from Eqs. (9) and (10) as

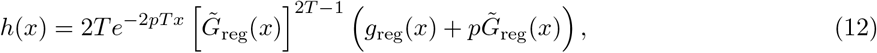

where 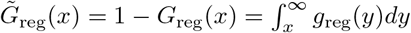.

### The length of a randomly chosen inter-crossover distance

Denote by *ϕ*(*x*) the density of a randomly chosen inter-crossover distance. Cox and Smith [46] proved a result on superposition of renewal processes that applies to our case. According to their Eq. (31), if the density of the distance to the next event (crossover) across all processes (meioses) is *h*(*x*), then the density of the length of a randomly chosen inter-crossover interval is given by

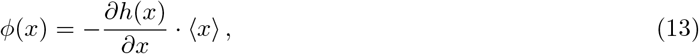

where 〈*x*〉 is the mean inter-crossover length (across all meioses). In our case, 〈*x*〉 = 1/(2*T*), and thus

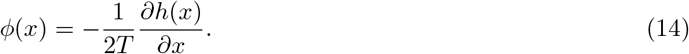

Substituting Eq. (12), and using the facts that 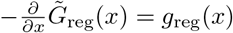 and 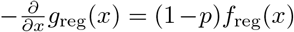,

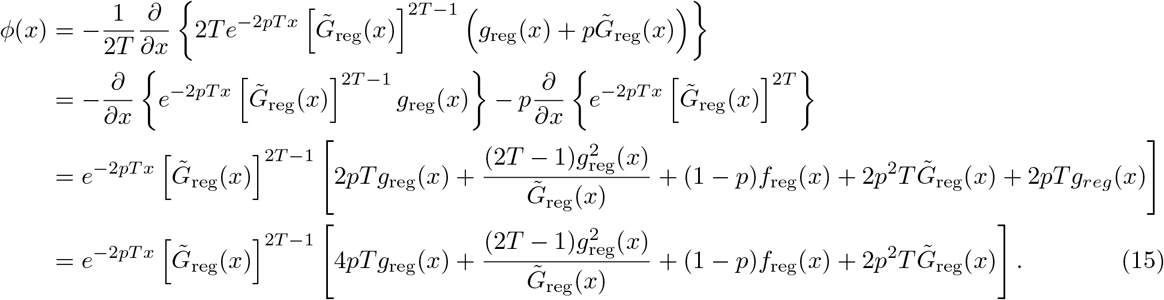

Eq. (15) is our main result, and is summarized as Eq. (1) in the main text. We note that as opposed to the Poisson model, inter-crossover distances under interference in a single meiosis are not independent, which is a general property of a superposition of renewal processes. To evaluate the various terms in Eq. (15) in our simulations, we truncated all sums at *k* = 50 and calculated all integrals using Matlab’s *integral* function.

### Finite chromosomes

The results above apply only to infinite-length chromosomes. To determine the distribution of segment lengths for finite chromosomes, we use a result derived by Gravel [47] in the context of local ancestry segments. Gravel showed that if a process along the chromosome partitions it into segments with a stationary length density *ϕ*(*x*), the density of segment lengths in a finite chromosome, *ϕ_L_*(*x*), is given by

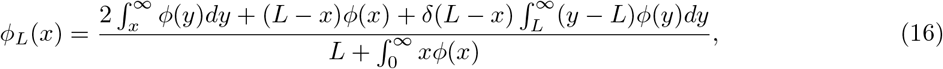

where *δ*(*L − x*) is the Dirac delta function, representing the probability that *x* spans the entire chromosome.

The theoretical distribution for the Poisson model, for an infinitely long chromosome, is *ϕ*(*x*) = 2*Te*^−2*Tx*^. Applying the finite chromosome correction of Eq. (16), we obtain

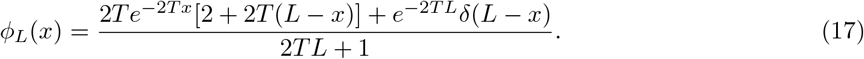

### Runtime analyses

To collect runtime statistics, we ran Ped-sim 1.0.1 and IBDsim 0.9-8 on a machine with four Xeon E5 4620 2.20GHz CPUs and 256 GB of RAM. We report wall clock time averaged from three runs of each program to produce IBD segments from 10,000 full siblings and 10,000 second cousins. To simulate the 10,000 full siblings with IBDsim, we used the following R code:

~~~
library(IBDsim)
quad <- nuclearPed(2)
res <- IBDsim(quad, sims=10000, query=list(‘atleast1’=3:4))
~~~

To simulate the siblings in Ped-sim, we used the following `def` file:

~~~
def full-sibs 10000 2
2 1
~~~

IBDsim defaults to assigning all non-founders as male and their spouses as female. We simulated 5,000 second cousin pedigrees with the default non-founder sex assignment, and the other 5,000 with female non-founders using the following R code:

~~~
library(IBDsim)
second_cousin_nonfound_male <- cousinPed(2)
res_male <- IBDsim(second_cousin_nonfound_male, sims=5000, query=list(‘atleast1’=11:12))
second_cousin_nonfound_female <- swapSex(second_cousin_nonfound_male, c(3,4,7,8))
res_female <- IBDsim(second_cousin_nonfound_female, sims=5000,
                                   query=list(‘atleast1’=11:12))
~~~

For Ped-sim, we simulated the same second cousin pedigree structures with the `def` file:

~~~
def second-cous-male 5000 4 M
4 1
def second-cous-female 5000 4 F
4 1
~~~

To benchmark Ped-sim’s time to produce genetic data for 4,450 full sibling pairs, we used input haplotypes from 8,955 individuals. These data are a subset of multiple sclerosis case-control samples [48] previously used to evaluate a multi-way relatedness inference method. Filters and phasing procedures used to generate this dataset are in the corresponding paper [6].

## Data availability

The SAMAFS genotype data are available from dbGaP under accessions phs000333.v1.p1 and phs000462.v1.p1. The Hemani *et al*. IBD proportions [34] are available from http://cnsgenomics.com/data.html#hemani. Pairwise IBD estimates for the SAMAFS relatives used in this study are available in S1 Dataset.

## Ethics statement

This study makes use of deidentified individuals from the SAMAFS and received exemption (#4) from IRB review from the Cornell University IRB (protocol 1408004874).

## Acknowledgments

We thank the reviewers for their helpful feedback and Monica Ramstetter for testing Ped-sim. We are grateful to the participants in the SAMAFS who made possible the real data evaluations. This study makes use of data generated by the Wellcome Trust Case-Control Consortium. A full list of the investigators who contributed to the generation of the data is available from www.wtccc.org.uk.

## Supporting information captions

**S1 Fig. Mean, Minimum, Median, and Maximum IBD sharing fractions in real and simulated data for full siblings through second cousins.** Points are from the SAMAFS, SAMAFS-validated subset (except full siblings), Hemani20k set (only full siblings), and the simulation models. The latter are labeled using abbreviations given in the main text. The SAMAFS and SAMAFS-validated values are mean-shifted for the first cousins, first cousins once removed, and second cousins, but are unaltered for the full sibling and the full sibling IBD2 quantities. Bars indicate one standard error as calculated from 1,000 bootstrap samples.

**S2 Fig. First cousins simulated using sex-specific maps have visually similar distributions of IBD sharing proportions relative to those simulated under a sex averaged map.** Sex-specific and sex averaged distributions heavily overlap both when using an interference (left) and a Poisson (right) model for inter-crossover distances.

**S3 Fig. Distributions of IBD sharing fractions for simulated full siblings and the real SAMAFS and Hemani20k full siblings.** Each simulation includes 10,000 full sibling pairs, the SAMAFS data are from the final 1,114 pairs, and 20,240 Hemani20k pairs.

**S4 Fig. Overlay of SAMAFS full sibling IBD distribution with that of simulated full siblings from each crossover model.** Plots are histograms of the 1,114 SAMAFS pairs and 10,000 simulated pairs generated under each of the crossover models, as indicated.

**S5 Fig. Overlay of Hemani20k full sibling IBD distribution with that of simulated full siblings from each crossover model.** Plots are histograms of the 20,240 Hemani20k pairs and 10,000 simulated pairs generated under each of the crossover models, as indicated.

**S6 Fig. Rates of inferring real and simulated relatives to their true degree of relatedness.** Degrees are inferred from kinship coefficients, with the latter calculated using inferred (for SAMAFS and SAMAFS-validated) or true (for the simulations) IBD segments (see Methods). Bars indicate one standard error based on 1,000 bootstrap samples over relative pairs.

**S7 Fig. Example pedigree structures for SAMAFS-validated relatives and for relatives descended from female-only non-founders.** (A) SAMAFS-validated pairs are required to be descended from a genotyped (black) full sibling pair and to have genotyped parent-child relatives that directly connect them to the full siblings. We further require that both the ancestral full sibling pair and all parent-child pairs be inferred as first degree relatives by Refined IBD. (B) Plot of female-lineage fourth cousins.

**S8 Fig. Number of IBD segments shared between maternal and paternal half-siblings.** (A) 10,0 simulated pairs for both types of half-siblings under the SS+intf model. (B) Maternal and paternal half-siblings within SAMAFS.

**S9 Fig. Local ancestry tract length distributions from simulations of 10,000 admixed samples.** Rates are from exponential fits (accounting for finite chromosomes; Methods) for each simulation scenario.

**S10 Fig. The effect of sex-specific maps on IBD segment lengths.** We used Ped-sim to simulate half-cousins with a common ancestor *T* = 1, 2, 4, 6 generations ago under the SS+Poiss model, extracting IBD segment lengths in bp for chromosome 1 (panels A-D, respectively). Each panel shows the simulated distribution of IBD segment lengths (over 10^5^ pairs for *T*= 1, 2 and 10^6^ pairs otherwise; purple circles), the theory from Eq. (2) (blue lines; includes the finite-chromosome correction of Eq. (16)), and the expectation based on a sex-averaged maps (red dashed lines). To evaluate Eq. (2) we replaced the integrals with sums over discrete coordinates, evenly separated by 10^4^ bp.

**S1 Dataset. Summary statistics and pairwise IBD proportions used to produce Fig 1, S1, S3, S4, S6, and results given in the text.**

