## Supplementary Figures for "Crossover interference and sex-specific genetic maps shape identical by descent sharing in close relatives"

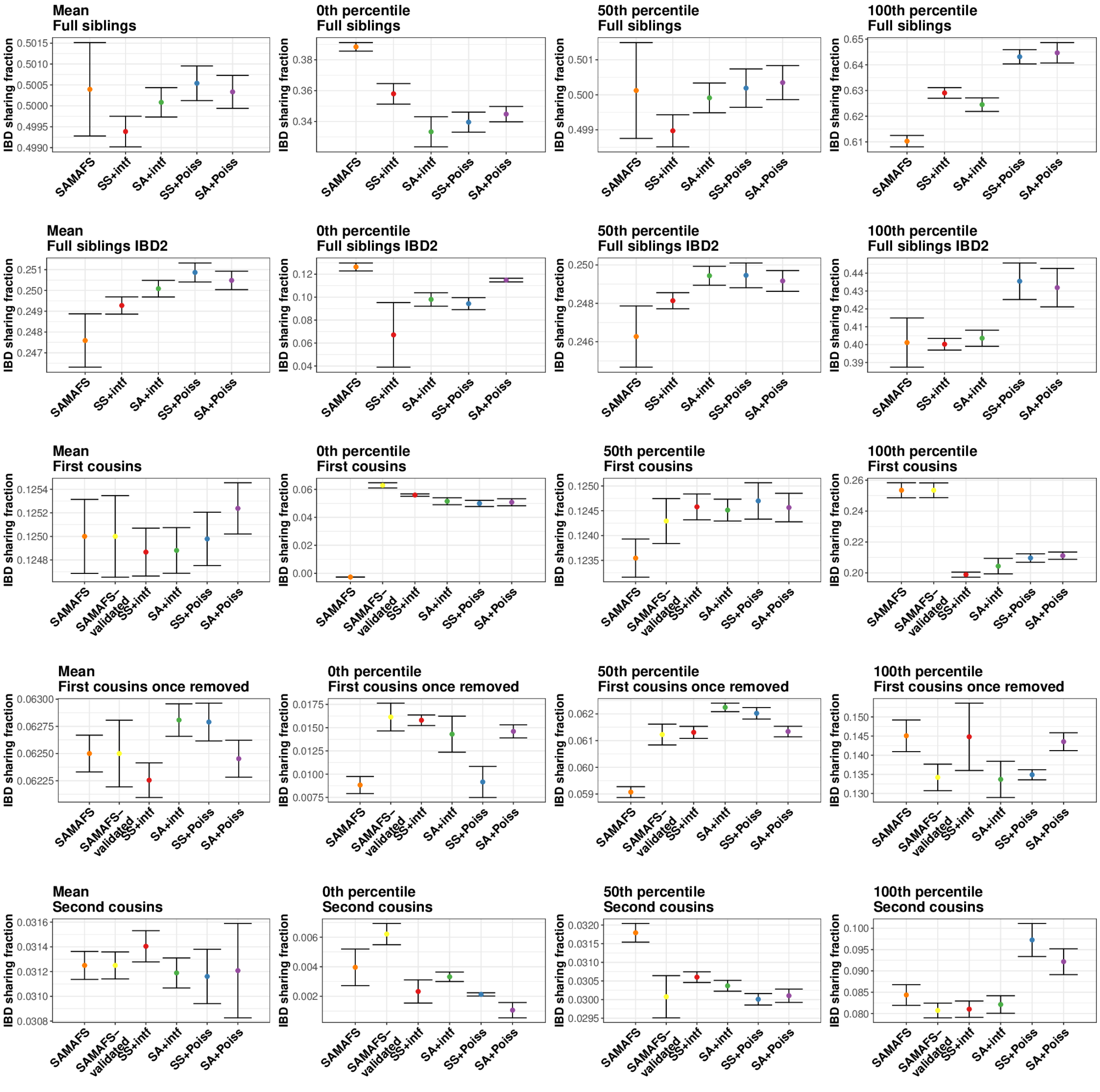


**S1 Fig: Mean, Minimum, Median, and Maximum IBD sharing fractions in real and simulated data for full siblings through second cousins.** Points are from the SAMAFS, SAMAFS-validated subset (except full siblings), Hemani20k set (only full siblings), and the simulation models. The latter are labeled using abbreviations given in the main text. The SAMAFS and SAMAFS-validated values are mean-shifted for the first cousins, first cousins once removed, and second cousins, but are unaltered for the full sibling and the full sibling IBD2 quantities. Bars indicate one standard error as calculated from 1,000 bootstrap samples.

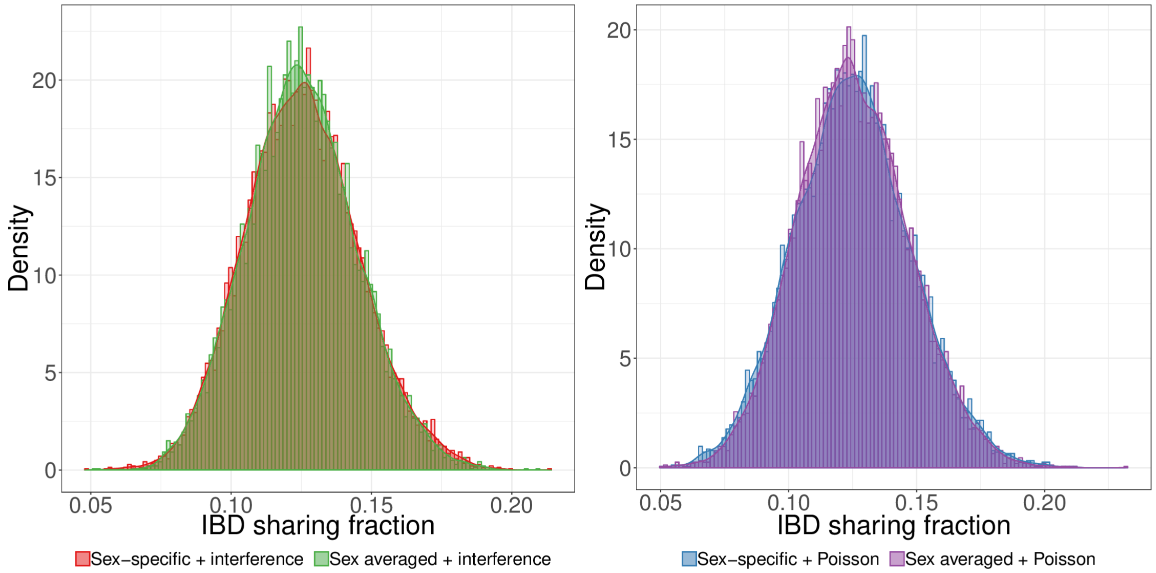


**S2 Fig: First cousins simulated using sex-specific maps have visually similar distributions of IBD sharing proportions relative to those simulated under a sex averaged map.** Sex-specific and sex averaged distributions heavily overlap both when using an interference (left) and a Poisson (right) model for inter-crossover distances.

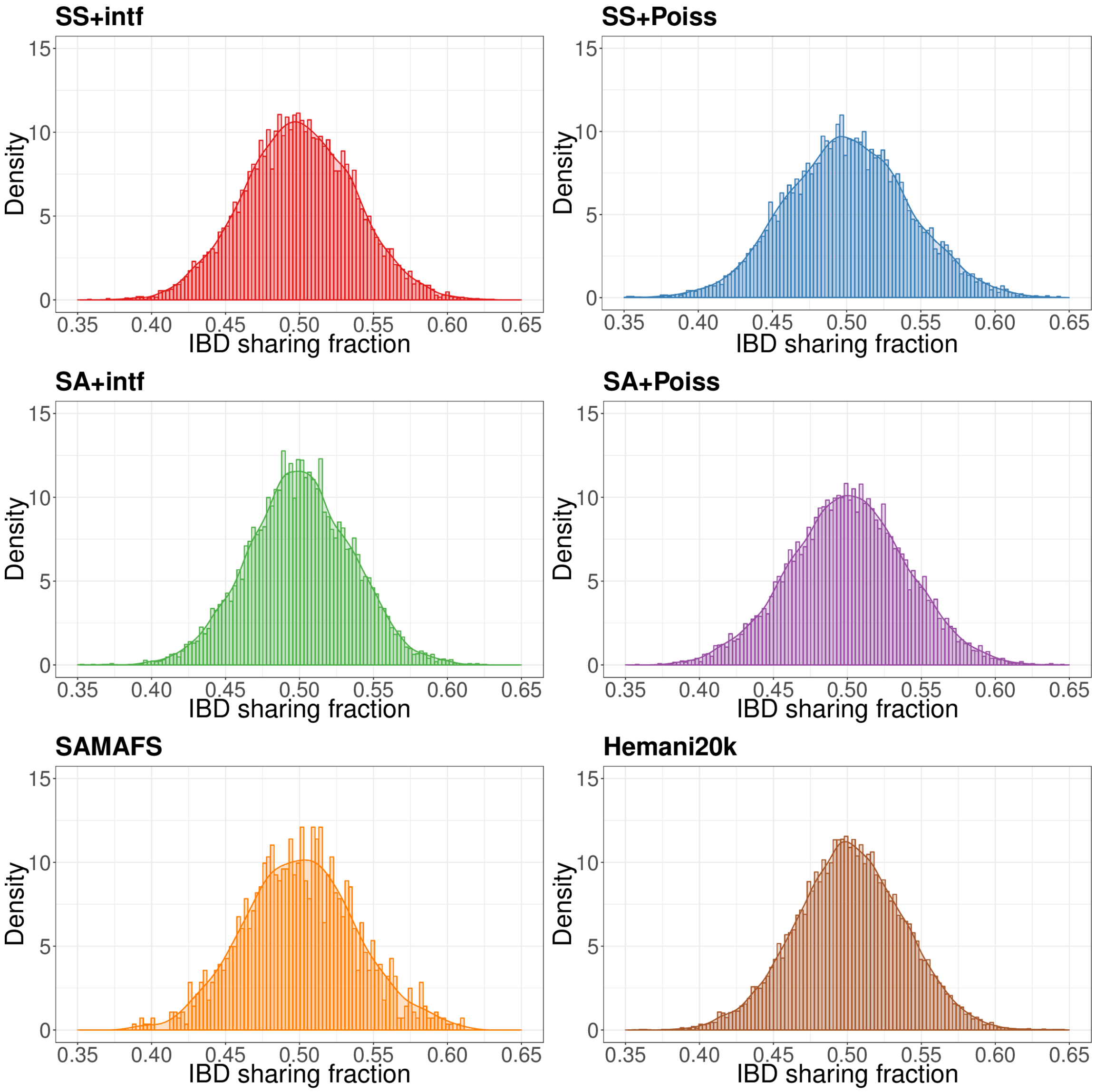


**S3 Fig: Distributions of IBD sharing fractions for simulated full siblings and the real SAMAFS and Hemani20k full siblings.** Each simulation includes 10,000 full sibling pairs, the SAMAFS data are from the final 1,114 pairs, and 20,240 Hemani20k pairs.

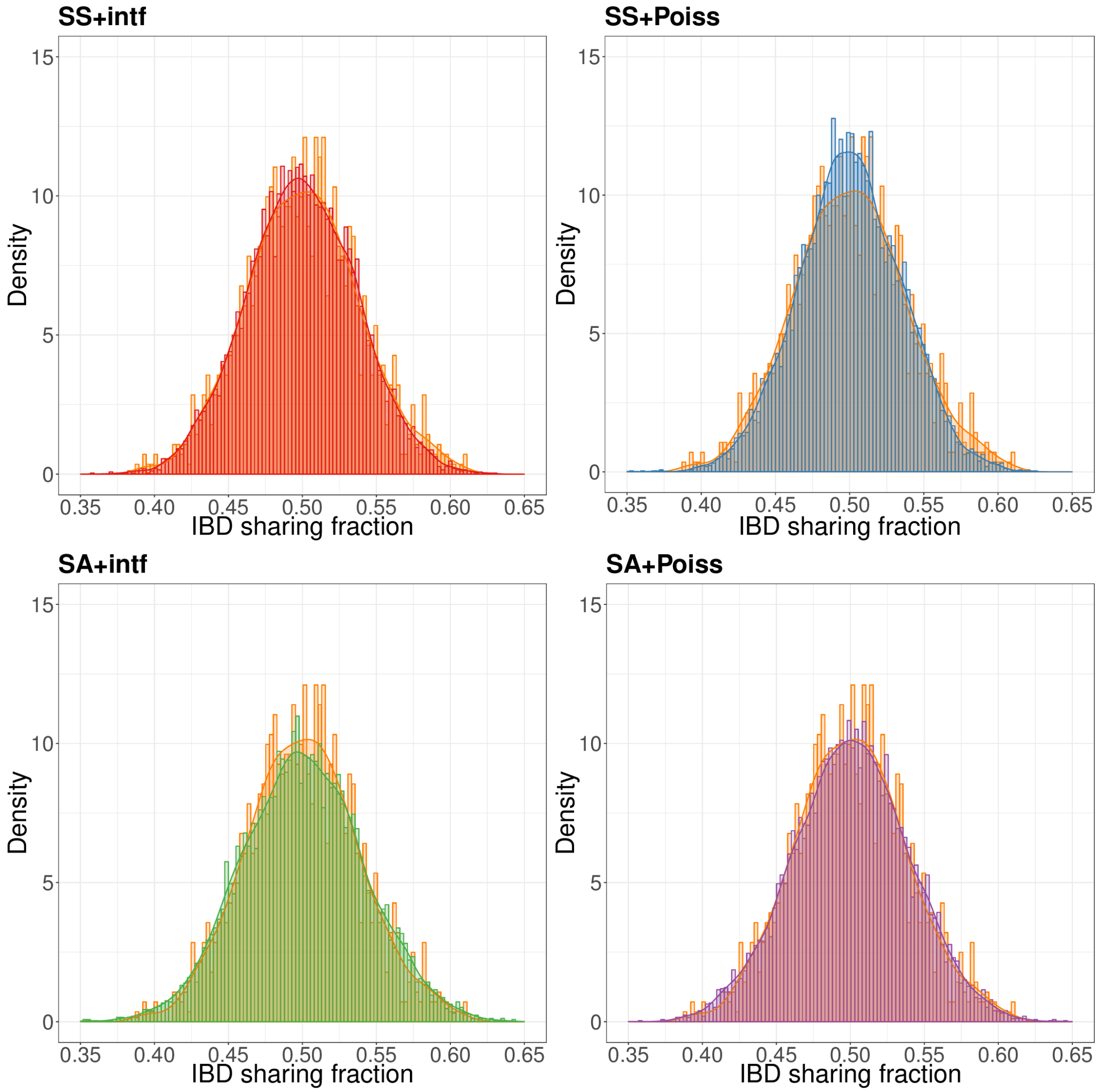


**S4 Fig: Overlay of SAMAFS full sibling IBD distribution with that of simulated full siblings from each crossover model.** Plots are histograms of the 1,114 SAMAFS pairs and 10,000 simulated pairs generated under each of the crossover models, as indicated.

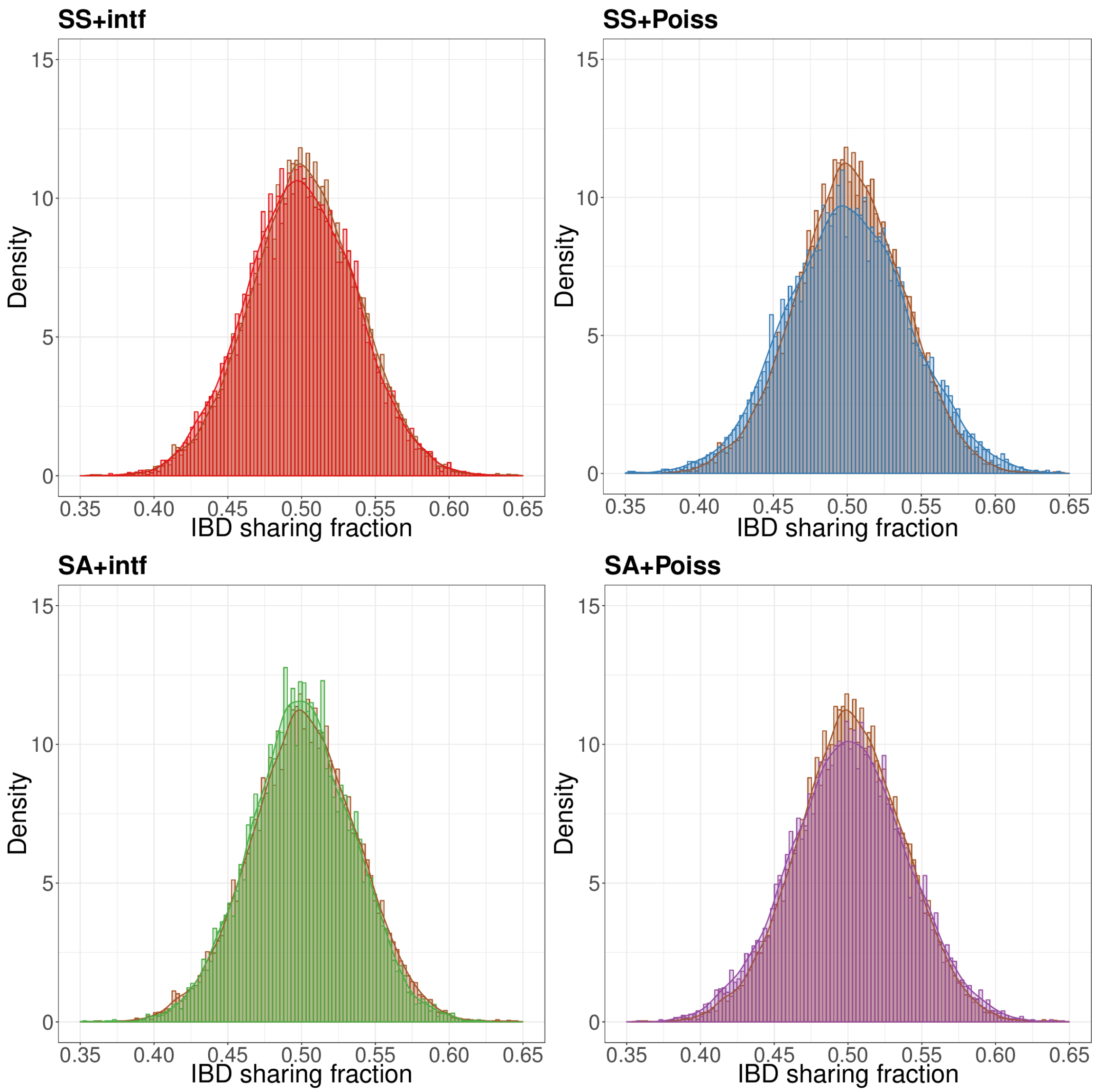


**S5 Fig: Overlay of Hemani20k full sibling IBD distribution with that of simulated full siblings from each crossover model.** Plots are histograms of the 20,240 Hemani20k pairs and 10,000 simulated pairs generated under each of the crossover models, as indicated.

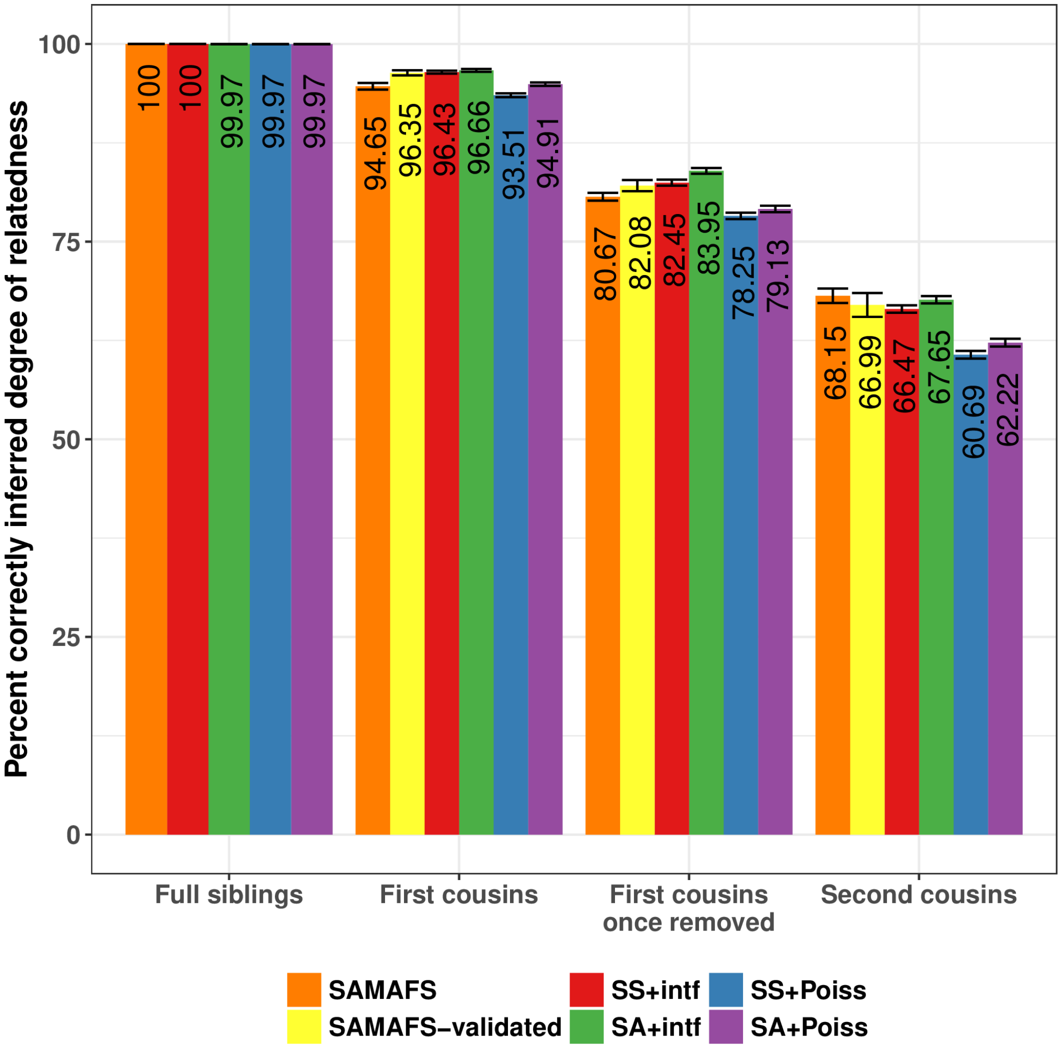


**S6 Fig: Rates of inferring real and simulated relatives to their true degree of relatedness.** Degrees are inferred from kinship coefficients, with the latter calculated using inferred (for SAMAFS and SAMAFS-validated) or true (for the simulations) IBD segments (see Methods). Bars indicate one standard error based on 1,000 bootstrap samples over relative pairs.

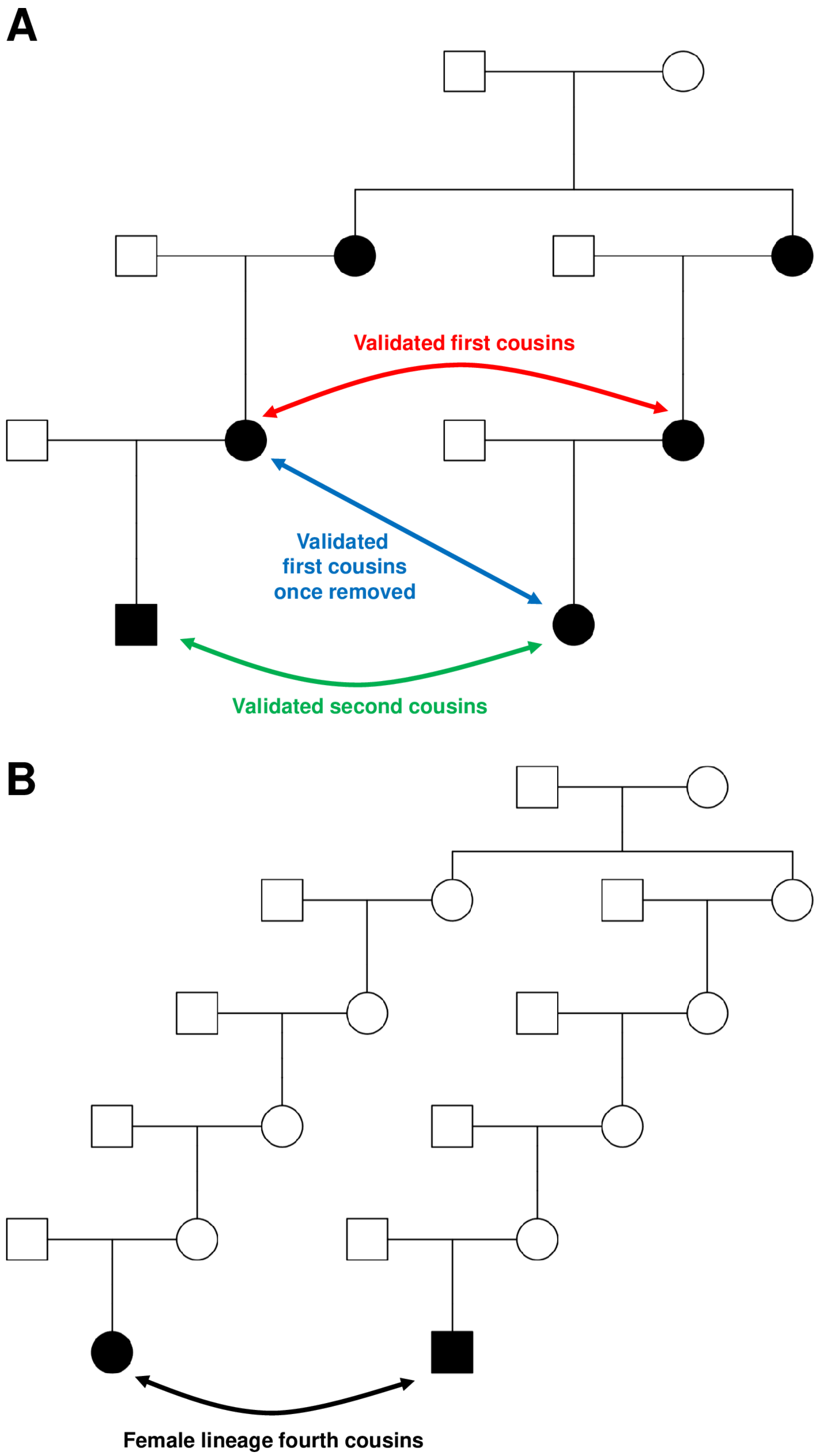


**S7 Fig: Example pedigree structures for SAMAFS-validated relatives and for relatives descended from female-only non-founders.** (A) SAMAFS-validated pairs are required to be descended from a genotyped (black) full sibling pair and to have genotyped parent-child relatives that directly connect them to the full siblings. We further require that both the ancestral full sibling pair and all parent-child pairs be inferred as first degree relatives by Refined IBD. (B) Plot of female-lineage fourth cousins.

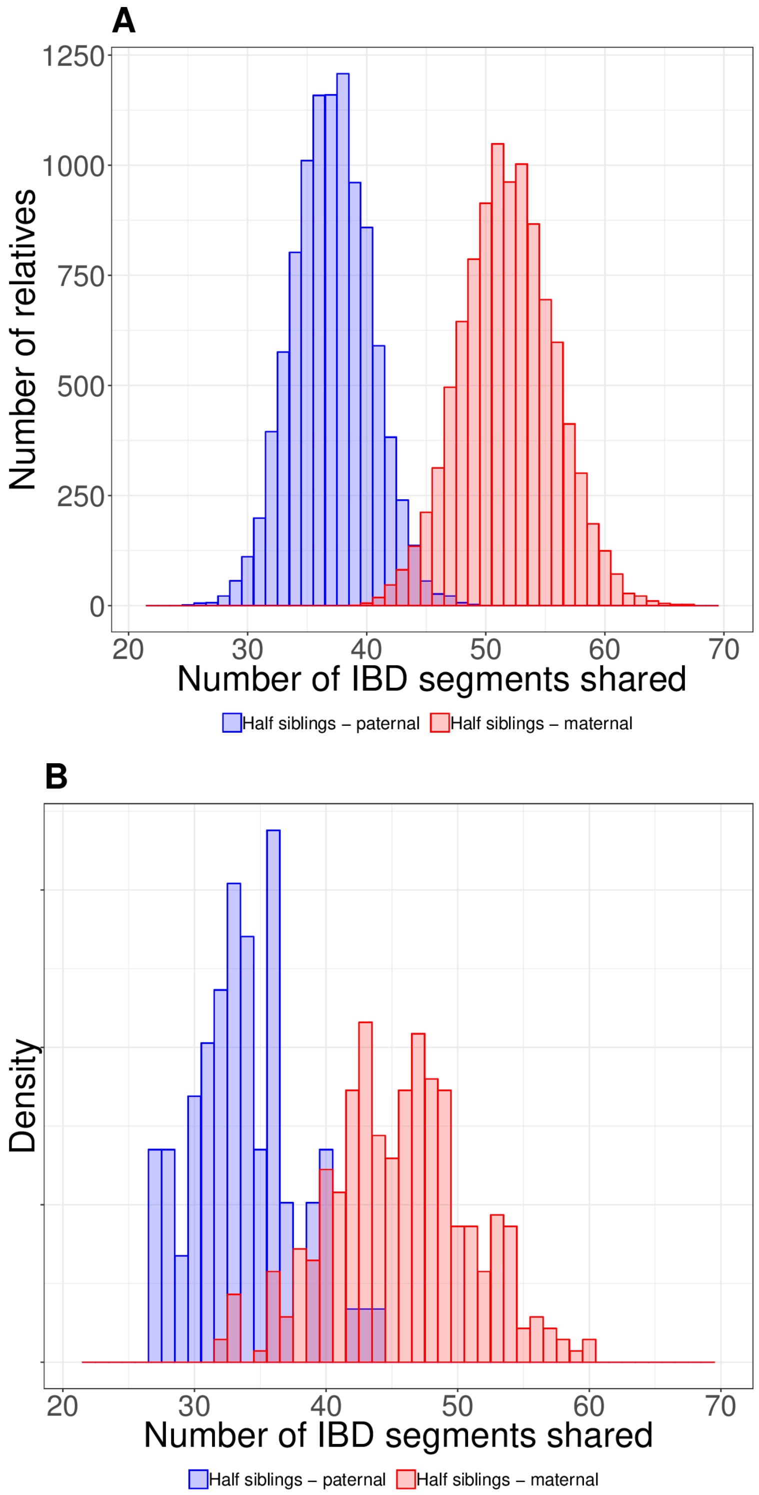


**S8 Fig: Number of IBD segments shared between maternal and paternal half-siblings.** (A) 10,000 simulated pairs for both types of half-siblings under the SS+intf model. (B) Maternal and paternal half-siblings within SAMAFS.

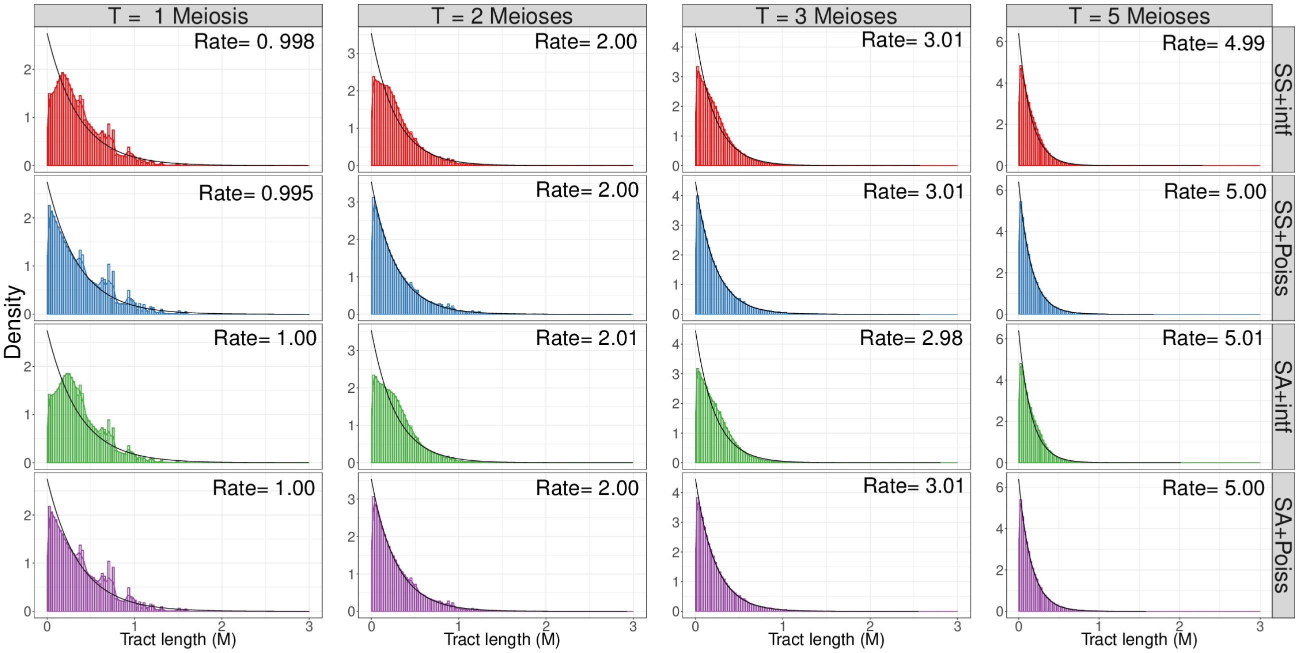


**S9 Fig: Local ancestry tract length distributions from simulations of 10,000 admixed samples.** Rates are from exponential fits (accounting for finite chromosomes; Methods) for each simulation scenario.

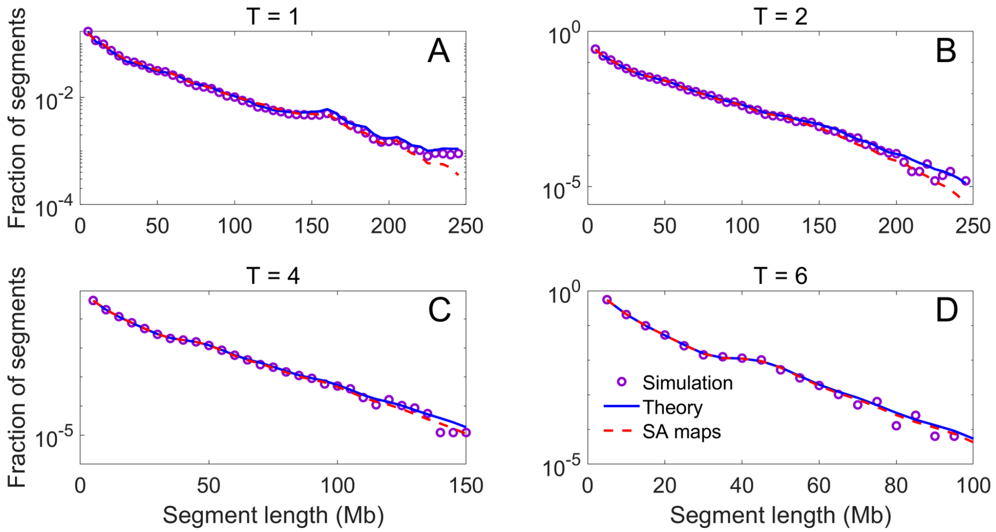


**S10 Fig: The effect of sex-specific maps on IBD segment lengths.** We used Ped-sim to simulate half-cousins with a common ancestor $T = 1, 2, 4, 6$ generations ago under the SS+Poiss model, extracting IBD segment lengths in bp for chromosome 1 (panels A-D, respectively). Each panel shows the simulated distribution of IBD segment lengths (over $10^5$ pairs for $T = 1, 2$ and $10^6$ pairs otherwise; purple circles), the theory from Eq. (2) (blue lines; includes the finite-chromosome correction of Eq. (16)), and the expectation based on a sex-averaged maps (red dashed lines). To evaluate Eq. (2) we replaced the integrals with sums over discrete coordinates, evenly separated by $10^4$ bp.
